# An *In-Silico* Investigation of Menthol Metabolism

**DOI:** 10.1101/618975

**Authors:** Taweetham Limpanuparb, Wanutcha Lorpaiboon, Kridtin Chinsukserm

## Abstract

Prevalence of mentholated products for consumption has brought great importance to studies on menthol’s metabolic pathways to ensure safety, design more potent derivatives, and identify therapeutic benefits. Proposed pathways of (-)-menthol metabolism based on metabolites found experimentally in previous works by Yamaguchi, Caldwell & Farmer, Madyastha & Srivatsan and Hiki et al. were not in agreement. This *in silico* approach is based on the three *in vivo* studies and aims to resolve the discrepancies. Reactions in the pathways are conjugation with glucuronic acid/sulfate, oxidation to alcohol, aldehyde & carboxylic acid, and formation of a four-membered/five-membered ring. Gas-phase structures, standard Gibbs energies and SMD solvation energies at B3LYP/6-311++G(d,p) level were obtained for 102 compounds in the pathways. This study provides a more complete picture of menthol metabolism by combining information from three experimental studies and filling missing links in previously published pathways.

## Introduction

(-)-Menthol or 1*S*,3*R*,4*S*-menthol is a naturally occurring compound found in plants of the *Mentha* genus commonly known as mint. It is the most abundant in nature among the 8 possible stereoisomers, and make up at least 50% of peppermint (*Mentha piperita*) oil and 70-80% of corn mint (*Mentha arvensis*) oil [1]. (-)-Menthol, commonly referred to as menthol, has characteristic minty smell and flavor and exerts a cooling sensation when applied to the skin and mucosal membranes [2]. Other isomers differ slightly in odor and physical characteristics and do not possess the cooling action [3, 4].

Menthol finds a wide range of applications from personal care products, medications, and confectionery to pesticides and cigarettes. The popularity of the compound as a flavoring agent ranks third most important after vanilla and citrus [5], and the annual production of menthol in India alone is in excess of 200 thousand metric tons [6]. Mentholated products can be purchased as prescribed or over-the-counter medications as alleviators of common cold and respiratory conditions [7], inhibitors of growth of foodborne pathogens [8], and analgesics [9].

Considering its wide range of applications, mechanisms of action of menthol were relatively unknown until recently. The cooling sensation is a result of the activation of transient receptor potential melastatin-8 (TRPM8), an ion channel selective to temperature, voltage, and menthol [10]. Experimental evidence also show that (-)-menthol can selectively activate κ-opioid receptors in mice and, as a result, leads to its analgesic properties [9]. In addition, chemical derivatives of menthol with enhanced activity have been successfully synthesized [11]. However, health effects of mentholated cigarettes is of great concern, not only because the improved taste may facilitate initiation or inhibit quitting but also because metabolism of menthol via this route of administration has not been well studied [12, 13].

A few studies have been conducted on toxicological effects of menthol which supports the generally accepted belief that it is safe and nontoxic. No signs of toxicity were observed in rats exposed to continuous doses of up to 800 mg/kg/day for 28 days [5], and chronic exposure to high concentrations of menthol vapor was not reported to have toxic effects in rats [14]. *In vitro* studies on various animal tissues report deterioration of biological membranes at concentrations 0.32-0.76 mM [15]. The recommended daily intake for humans of 0-0.2 mg/kg proposed by the WHO [16] is not supported by any toxicological data but was set to err on the side of safety knowing that higher doses taken may not have produced adverse side effects.

To the best of our knowledge, three *in vivo* studies by Yamaguchi, Caldwell & Farmer [17], Madyastha & Srivatsan [18] and Hiki et al. [19] have identified metabolites of menthol in humans and animals. Metabolites were identified by GC/MS from the urine and bile of rats treated with oral doses of 500 [17] and 800 [18] mg menthol/kg body weight. Over the course of 48 hours, a majority of the doses were excreted in the urine and feces. A more recent randomized, double-blind, placebo-controlled study in human by Hiki et al. [19] was conducted by directly spraying 0.8% (-)-menthol solution at escalating doses of 10-40 mL onto the gastric mucosa. Blood and urine of the participants were sampled over a 24-hour period and analyzed with GC/MS for menthol metabolites. In total, 72 metabolites were identified or proposed in this human study alone, compared to 9 and 18 metabolites in the previous two experiments. (See S3 File for the full list of metabolites in the first worksheet in the spreadsheet file.) *In vitro* investigation of metabolism in human liver microsomes revealed that the same key reactions in the metabolic pathway in rats occur in the microsomes [20, 21].

This *in silico* investigation is based on the metabolites identified experimentally by the three *in vivo* studies [17-19]. We aim to resolve discrepancies and missing links found in these three studies by proposing more complete pathways in Figure 1 where all 73 experimentally identified metabolites, 5 previously proposed intermediates and 24 newly proposed intermediates are included. Possible reactions involved in the pathways are conjugation with glucuronic acid/sulfate, oxidation to alcohol, aldehyde & carboxylic acid, and formation of a four/five-membered ring at position 3, 7, 8, 9 and 10 of the parent compound. In this paper, we calculated Gibbs energies of reactions and associated them with the type, the position and the step of reaction in the pathways.

**FIGURE 1.**
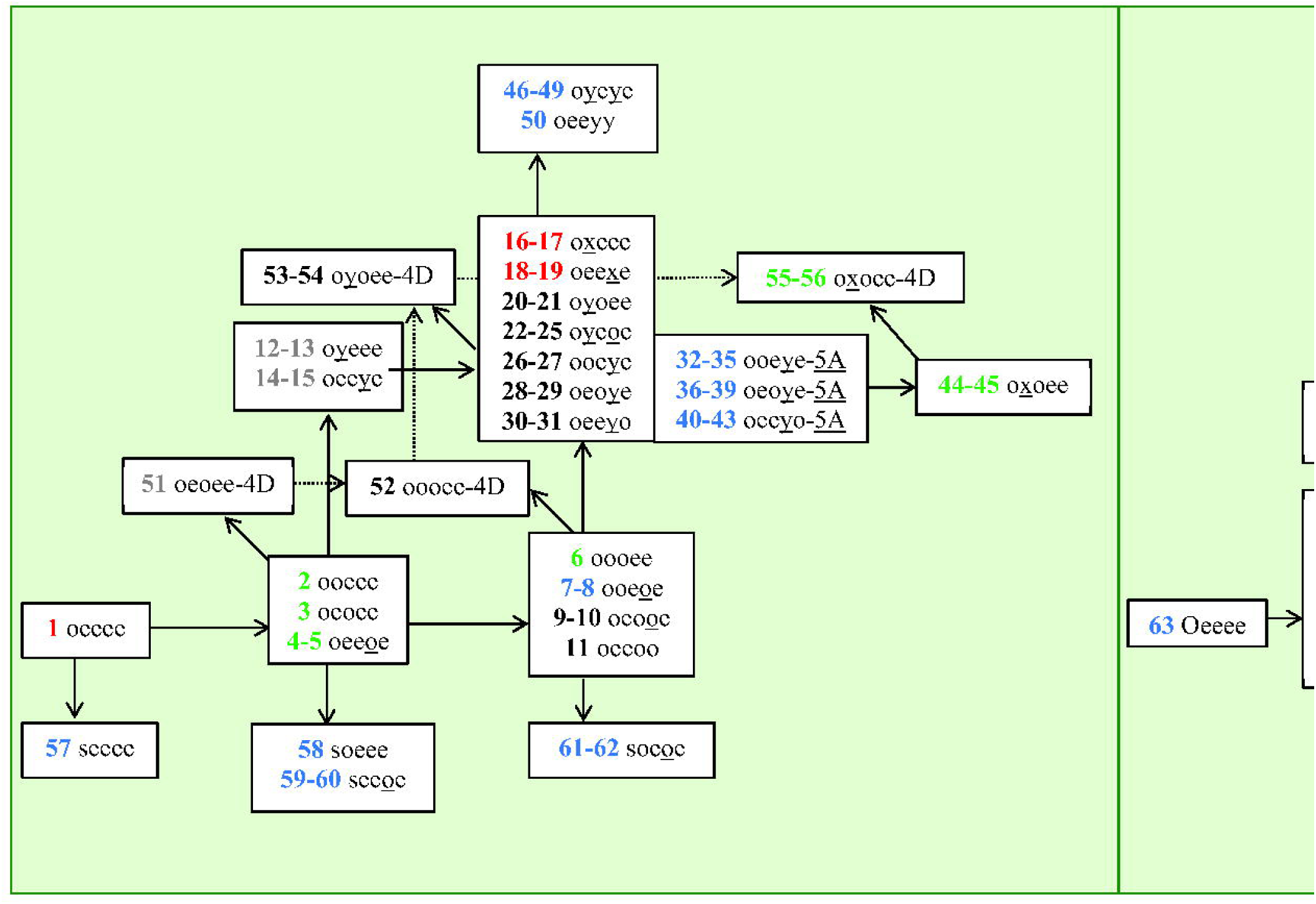
Metabolic pathway of menthol in rats and in human, an adaptation from Yamaguchi, Caldwell, & Farmer [17], Madyastha & Sirvastan [18] and Hiki et al. [19]. Red, Green, and Blue numbers indicate that menthol metabolites were found in both rats and human, only in rats, and only in human respectively. Gray and Black numbers indicate menthol metabolites proposed by previous experiments and by this paper respectively. Arrows to the right and arrows upward indicate oxidation reactions 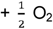 and – H_2_ respectively. Downward arrows indicate conjugation with sulfate. Dashed arrows indicate reactions of four-membered ring metabolites. Diagonal arrows toward top left indicate dehydration reaction. Main pathways are shown on the left and pathways containing glucuronide metabolites with similar possible connections are shown on the right. Lists of compounds and reactions are provided in Table 1 and Table 2, respectively.

### Computational Details

Gas-phase structures were calculated at the B3LYP/6-311++G(d,p) level of theory and were confirmed to be at minimum energy on the electronic potential energy surfaces by frequency calculations. The solvation energies in water of the gas-phase structures were calculated with the SMD model [22] through single-point energy calculations. The calculation of Gibbs energies in solution phase is the same as in our previous work [23, 24] where there is a correction for the standard state of liquid water (∼ 55 mol/L) [25-28]. All quantum chemical calculations were performed using the Q-Chem 5.1 program package [29]. (Shell script, spreadsheet templates, and Mathematica [30] notebook used were modified from our previous work [23, 24]. All output files and other associated codes to obtain the standard Gibbs energies of the reaction are provided in S1 File and S2 File respectively.) The abbreviated names for each of the metabolites in this study are as in Table 1. For simplicity, we based the naming system of menthol metabolites on their five substitutable positions, namely position 3, 7, 8, 9 and 10. A menthol metabolite is referred to as a five-character sequence named according to its substituted functional groups at these positions with the abbreviation explained below in Table 1.

**TABLE 1.**
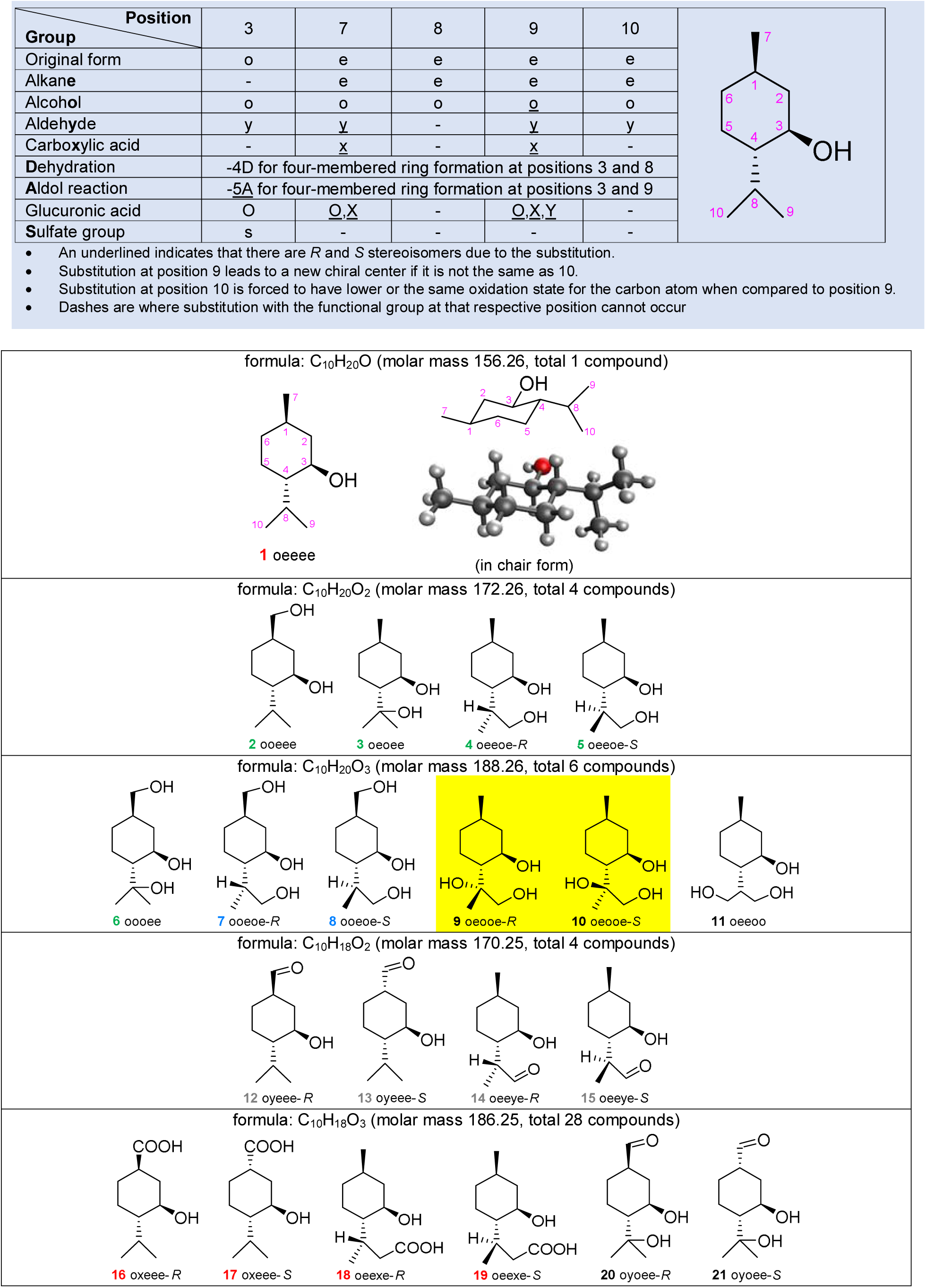
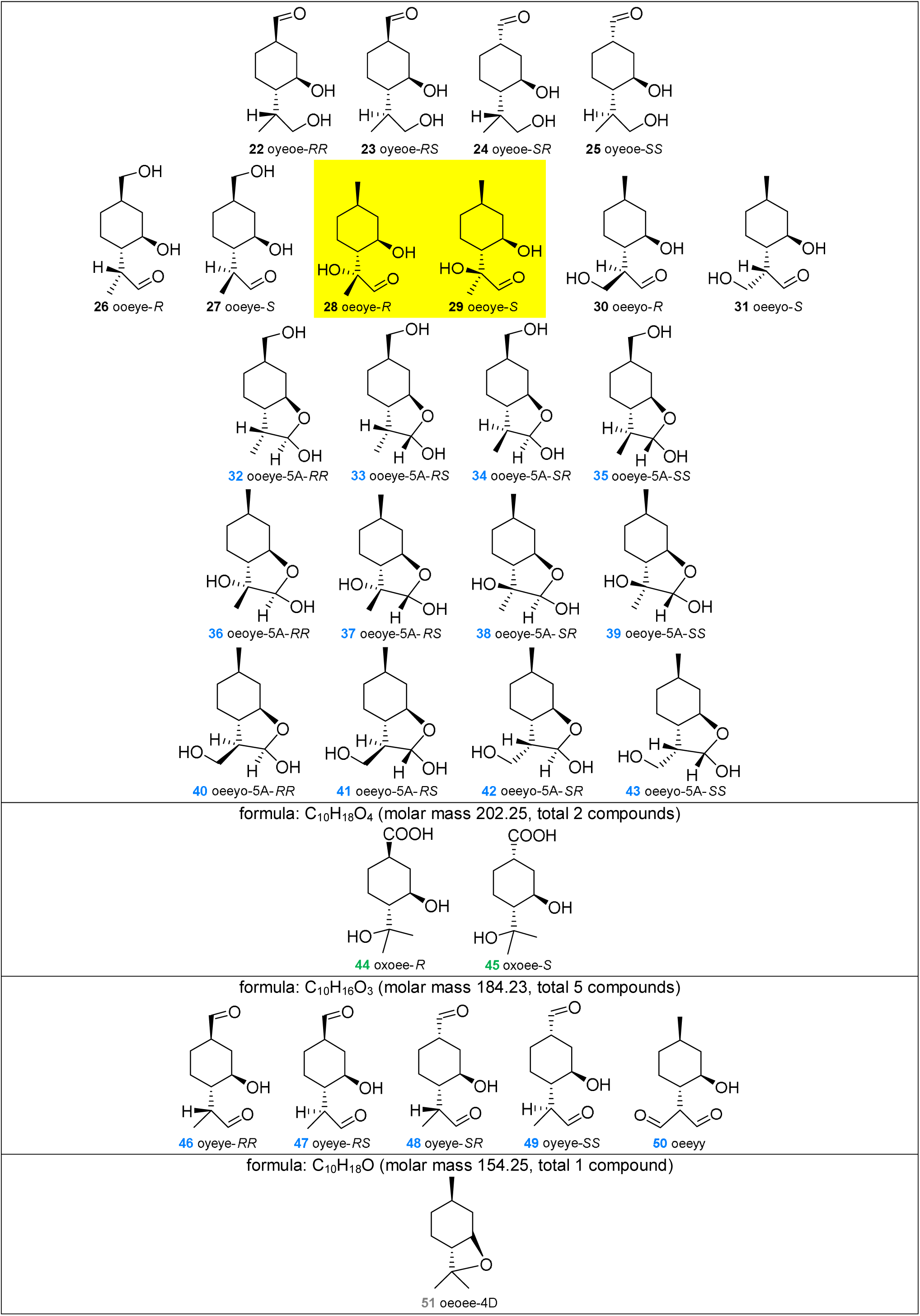
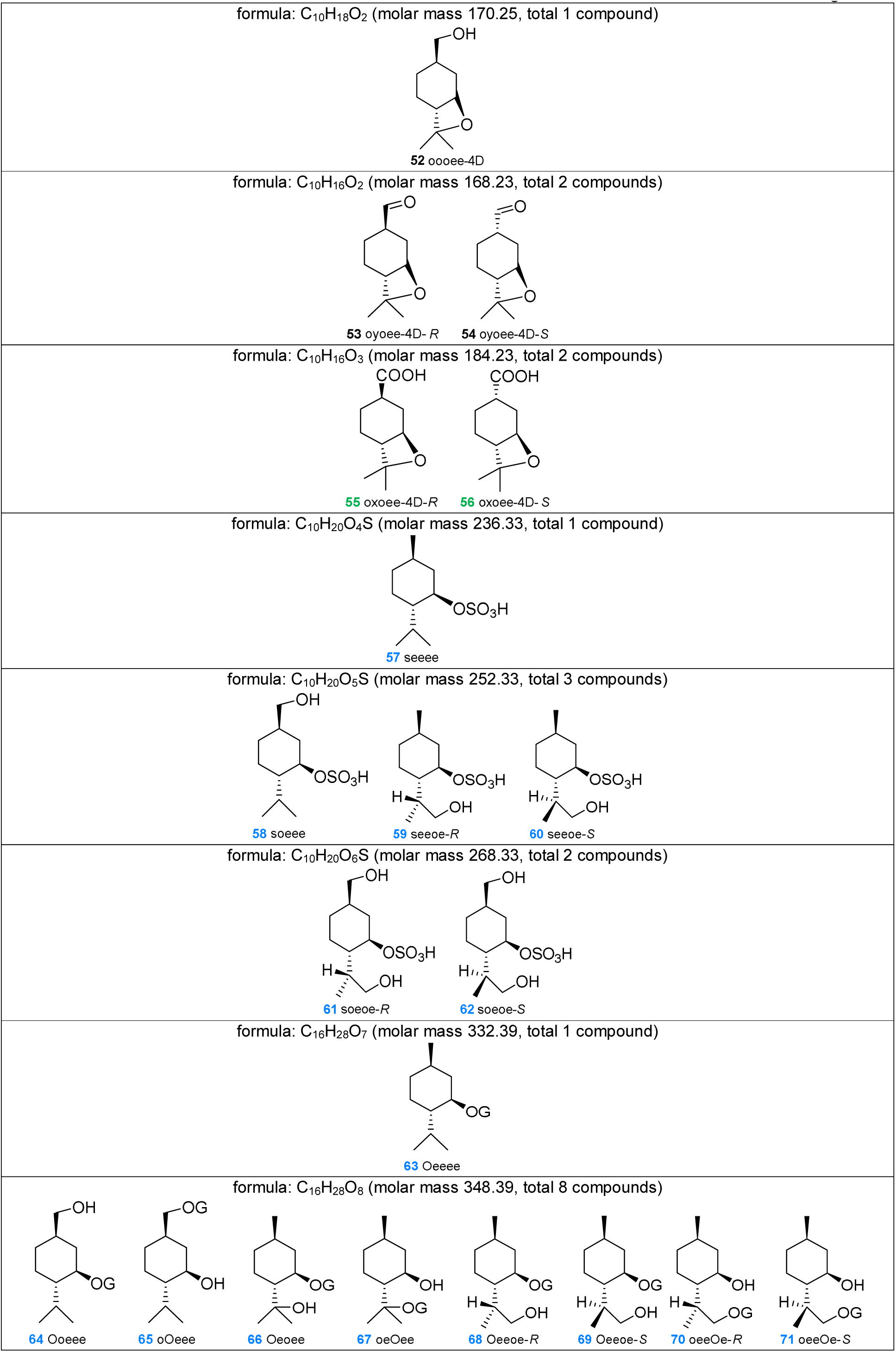
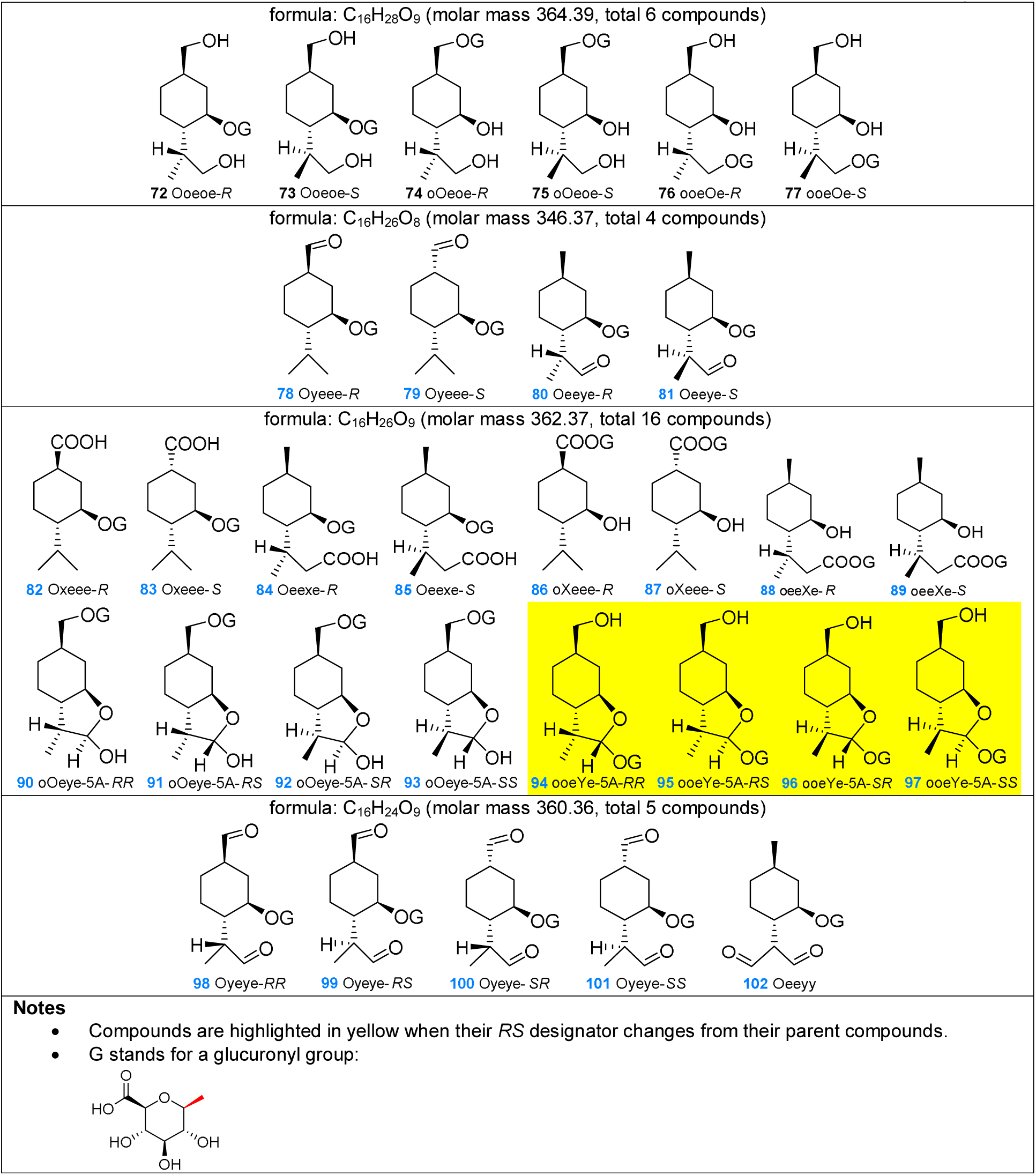
Abbreviations for the nomenclature of menthol metabolites referred to by the present study and a list of 102 compounds in this study grouped by molecular formula.

**TABLE 2.** Representative of oxidation reactions to alcohol, aldehyde and carboxylic acid, ring formation (dehydration reaction and aldol reaction) and conjugation with glucuronic acid/sulfate group.

| Abbreviation/explanation | Example |
| --- | --- |
| o1 for oxidation from alkane to alcohol |  |
| o2 for oxidation from aldehyde to carboxylic acid |  |
| o3 for oxidation from alcohol to aldehyde |  |
| 4D for dehydration (four-membered ring formation) |  |
| 5A for aldol reaction (five-membered ring formation) |  |
| g for conjugation with glucuronic acid |  |
| s for conjugation with sulfate |  |

All DFT calculations were completed with no imaginary frequencies, showing that each of the structures obtained from gas-phase calculations were minima on the potential energy surfaces. The lowest energy structure of (-)-menthol is a chair conformer of hexane where all three substituent groups are in equatorial positions as shown in Table 1. This is consistent with previous computational result at B3LYP/6-31G(d,p) level [31]. Benchmark calculations were also performed at MP2/6-311++G(d,p) level for metabolites along the most likely pathways in Figure 2. Reaction energies obtained from MP2 and B3LYP are in good agreement. (Coefficient of determination *r*^2^=0.9999 and mean absolute error (MAE) of 2.25 kcal/mol. See the third worksheet of the Excel file in S3 File for details.) These results are in line with earlier studies and confirm that B3LYP/6-311++G** yields acceptable results at a reasonable computational cost [23, 32].

**FIGURE 2.**
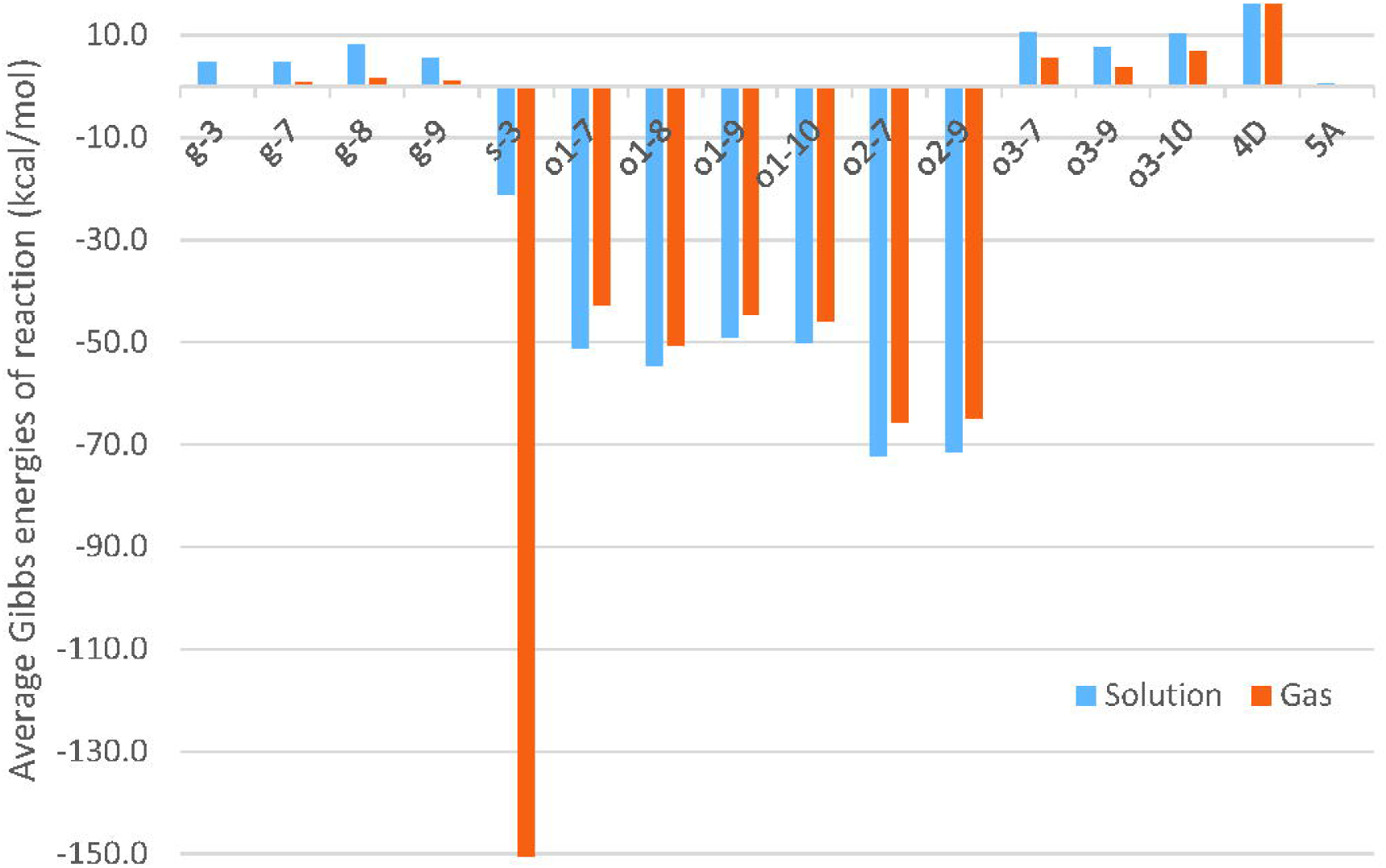
Average Gibbs energies of reaction in the gas phase and in solution for various reactions at the B3LYP/6-311++G(d,p) level of theory: (o1) oxidation from alkane to alcohol, (o2) oxidation from aldehyde to carboxylic acid, (o3) oxidation from alcohol to aldehyde, (4D) dehydration or four-membered ring formation, (5A) aldol reaction or five-membered ring formation, (g) conjugation with glucuronic acid and (s) conjugation with sulfate at five different positions of (-)-menthol. Representative reactions of each type are shown in Table 2.

## Results

The present study has combined the different published metabolic pathways of menthol and offers the relative stabilities of each metabolite based on thermodynamic calculations for each step involved as reported in Figures 1 to 4. Reaction energies were computed with the relevant additional reagents (oxygen, sulfate group, hydronium ion and glucuronic acid) and product (water and hydrogen) added to the scheme. They may not be the actual compound in the reactions but they serve as simple reference points for the thermodynamic calculations for reactions of interest. The full list of compounds and reaction energies is in the first worksheet of the Excel file in S3 File.

## Discussion

The standard Gibbs energies for each reaction in both gas and solution phase listed in the first worksheet of the spreadsheet of S3 File are summarized in Figure 2. With the exception of sulfation (s-3), reaction energies in gas phase are lower in magnitude but have the same sign as their equivalents in solution phase. Oxidation reactions (o1 and o2, addition of 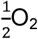) and conjugation with sulfate are the most exergonic and should occur easily. This may be explained by the fact that oxidation tends to introduce polar functional groups whose interactions with water serve to stabilize the compound. The reaction energies of sulfation are very exothermic in gas phase but not solution phase. This large difference in energy is explained by the presence of charged species in the reactant side which is greatly stabilized by the solvent – water – when compared to the product side that receives little stabilization upon solvation. The most endergonic reactions are four-membered ring formation (4D) and oxidation from alcohol to aldehyde (o3, removal of H_2_). The four-membered ring formation was proposed based on experimental evidence [18] published in 1988 and should be verified in further experiment. Difference in reaction energies due to position effect can be mostly explained by steric hindrance (i.e. g-8 has the highest reaction energy.) and inductive effect (i.e. o-8 producing secondary alcohol is the most exergonic.).

Gas phase energies were used to calculate solution phase energies, but as discussed previously they are not representative of the reactions that occur in biological systems. Since these reactions happen in solution, only the energies in solution phase were considered for each metabolite. Standard Gibbs energies for each metabolite are shown relative to the parent compound (**1** oeeee) in Figure 3. The step numbers correspond to the number of reactions required to generate each compound from **1** oeeee according to Figure 1. The first step from the parent compound tends to be the most exergonic with an average at −38.6 kcal/mol and the average reaction energy decreases monotonically to around −3.1 kcal/mol at the fifth step.

**FIGURE 3.**
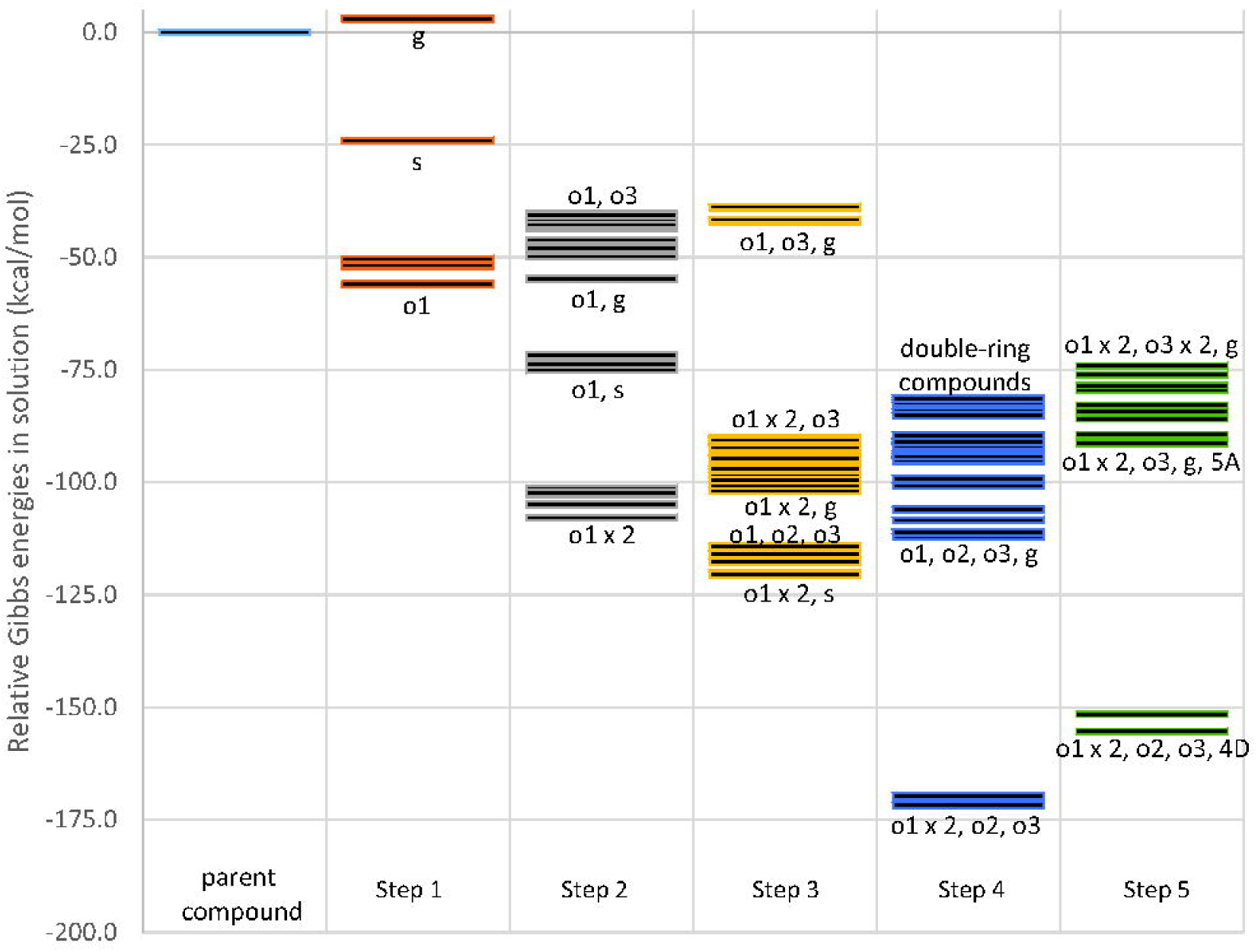
Relative stability of 102 (-)-menthol metabolite compounds at the B3LYP/6-311++G(d,p) level of theory.

Compounds with the lowest energy from each step were identified and are shown in Figure 4 with additional intermediates for completion. In general, the metabolite with the lowest relative energy in a step was the starting material for the lowest energy metabolite in the next.

**FIGURE 4.**
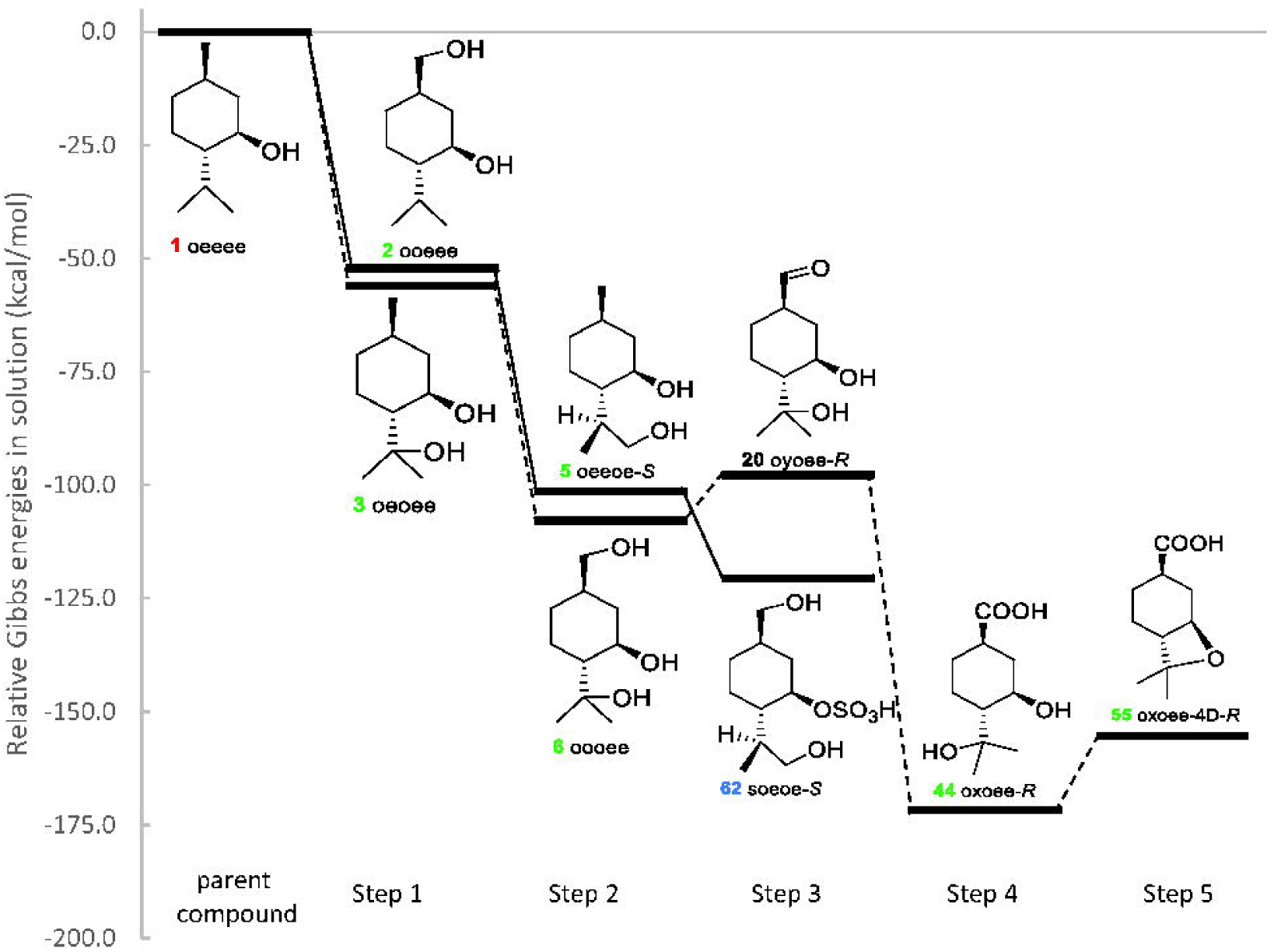
Lowest Gibbs energy diagram for the metabolism of (-) menthol at the B3LYP/6-311++G(d,p) level of theory.

- This energy diagram (Figure 4) is in agreement with major aspects of published metabolic pathways, in particular the conversion of menthol to *p*-menthane-3,8-diol (**3** oeoee). Partly due to increased solubility, the compounds **3** oeoee and its glucuronic acid conjugates, **66** Oeoee/**67** oeOee, were found to be major metabolites excreted in the urine of both rats and humans [17-19]. In contrast, *p*-menthane-3,7-diol (**2** ooeee) and *p*-menthane-3,9-diol (**4** oeeoe-*R*, **5** oeeoe-*S*) excreted from both rats and humans in small quantities. Figure 2 reports that oxidation from alkane to aldehyde at either position 7, 8, 9, or 10 is equally exothermic with a slight preference for position 8. Published evidence that **3** oeoee is formed as a product of enzymatic activity [18] and this observed thermodynamic preference explain the disproportionately large amount of **3** oeoee isolated experimentally compared to its isomers.
- Oxidation from alcohol to aldehyde is an endothermic reaction, hence metabolites containing aldehyde groups are either not detected or detected in small quantities and serve as intermediates to products of intramolecular aldol condensation to form cyclic ethers or further exothermic oxidation to carboxylic acid. In rats, no metabolites containing aldehyde groups were detected in the plasma, urine, bile, or feces. [17, 18] The published metabolic pathways show a direct conversion from alcohol to carboxylic acid. Only the most recent study conducted by Hiki et al. [19] reported detection of aldehyde menthol glucuronides in human urine at very low levels; the pathway proposed by Hiki et al shows further conversion to cyclic ethers and carboxylic acid. Since oxidation is a stepwise process, Figure 4 shows this stepwise conversion from **6** oooee to **44** oxoee-*R*.

### Concluding remarks

In this study, gas-phase structures of menthol and its metabolites (a total of 102 compounds and 151 reactions) were obtained by quantum calculations at B3LYP/6-311++G(d,p) level. The standard Gibbs energies of their respective reactions in solution were calculated with the SMD solvation model and corrected for standard state conditions. The lowest energy diagram (Figure 4) reported was largely in agreement with previously published experimental results. Information obtained in this study opens possibilities for further investigation of the pharmacological effects of menthol and its metabolites. Given that oxidation metabolites of menthol are energetically favorable, potency and toxicity of these oxidized derivatives should be further investigated. Different stereoisomer of menthol as well as MD-based approaches could also be explored in future research.

## Supporting information

S1 File All Q-CHEM output files (zip)

S2 File Wolfram Mathematica notebook, shell script and template (zip)

S3 File Microsoft Excel spreadsheet and associated files (zip)

## Acknowledgement

We appreciate helpful suggestions from Assoc. Prof. Dr. Yuthana Tantirungrotechai.

## Author Contributions

[This is to be copied from Editorial Manager after acceptance of the manuscript.]

