## Supplementary material for "An *In-Silico* Investigation of Menthol Metabolism": S1 File: README.txt

DOI: 10.1371/journal.pone.0216577

- File naming convention follows the paper and hence is case-sensitive.  This system is native in Linux but can be enable in other operating systems.  For Windows 10, please refer to Windows_10_case-sensitivity.txt for further instruction.
- All output files are generated by Q-Chem (www.q-chem.com) and can be opened in IQ-mol (iqmol.org).

- B3LYP folders contains all Q-Chem output files calculated at B3LYP/6-311++G**
  + o-series folder contains the parent compound and its oxidized products.
  + g-series folder contains all glucuronide metabolites
  + s-series folder contains the sulfate metabolites.
  + common folder contains all other necessary reactants and products.

- MP2 folders contains all Q-Chem output files calculated at MP2/6-311++G**

- In addition to output files, these files contain extracted information in each folder:
  + 0_file_list.txt is the list of all files.
  + 1_final_energy.csv contains the electronic energy. (in AU)
  + 2_solvation.csv contains the solvation energy. (in kcal/mol)
  + 3_zero_point.csv contains the zero-point energy (ZPE) correction. (in kcal/mol)
  + 4_enthalpy.csv contains the enthalpy correction. (in kcal/mol)
  + 5_entropy.csv contains the entropy. (in kcal/mol)
