## Supplementary material for "An *In-Silico* Investigation of Menthol Metabolism": S2 File: README.txt

DOI: 10.1371/journal.pone.0216577

- script.txt contains unix commands to generate input files, to run them and to extract information from them.
- template-B3LYP.txt contains Q-Chem instructions to optimize, calculate vibrational energy and solvation energy of a structure.
- calculations.nb is a Mathematica's notebook file that read Q-Chem output from "qchem output" folder and generate reaction energy output in "tables and figures" folder.
- list-reactions.txt contains all stepwise reactions for calculations.nb to read.
- list-reactions.txt contains all reactions starting from the parent compound for calculations.nb to read.
