## Supplementary material for "An *In-Silico* Investigation of Menthol Metabolism": S3 File: calculations.pdf

Compounds and reactions

| Calculation procedure | Name | Quantity | Unit | Ref |
|---|---|---|---|---|
| 1. Optimized in gas phase at B3LYP/6-311++G(d,p) | Hartree | 627.5094742 | kcal mol^-1/AU | 2014 CODATA |
| 2. Frequency calculation in gas phase at B3LYP/6-311++G(d,p) | Temp | 298.15 | K | Room Temp |
| 3. Single point energy calculation in water (SMD) at B3LYP/6-311++G(d,p) | Faraday constant | 96485.3329 | C/mol | 2014 CODATA |
| | calorie | 4.184 | J/cal | ISO 31-4:1992 |
| Q-CHEM Developer Version 5.1 | R | 8.3144598 | J mol^-1 K^-1 | 2014 CODATA |
| Program default output: STANDARD THERMODYNAMIC QUANTITIES AT 298.15 K AND 1.00 ATM | R | 0.082057338 | L atm mol^-1 K^-1 | 2014 CODATA |
| | [H2O] | 55.34 | M | Handbook of Chemistry and Physics, CRC press, 64th Ed |
| | ΔG o->* | 0.0030 | AU | Limpanuparb2017 Section 2.4 |
| | ΔG H2O | 0.0038 | AU | Limpanuparb2017 Section 2.4 |

The energies were adjusted to solution standard state conditions * – 1M, 298.15 K – with the exception of water where 55.34 M was used.

| Formula/Filename | Number in the paper | Other name | E+G/$C$1+C$9 | F+I/$C$1-$C$2*J/1000/$C$1 | Q-CHEM output at line: Final energy is | (9) = (6) - (0) | Zero point vibratio | Total Enthalpy: | Total Entropy: | # Imag | Ref | | | | Conclusion |
|---|---|---|---|---|---|---|---|---|---|---|---|---|---|---|---|
| | | | $G^*_{soln}$ | $G_{gas}$ | $E_{elec}$ | $\Delta G_{solv}$ | ZPE | $H_{corr}$ | S | | 17 | 18 | 19 | This paper | |
| | | | AU | AU | AU | kcal/mol | kcal/mol | kcal/mol | cal/mol/K | | | | | | |
| C6H10O7 | | | -761.357 | -761.313 | -761.452 | -29.6 | 111.6 | 120.4 | 112.2 | 0 | | | | | |
| H2 | | | -1.175 | -1.181 | -1.180 | 1.6 | 6.3 | 8.4 | 31.1 | 0 | | | | | |
| H2O | | | -76.462 | -76.455 | -76.459 | -8.6 | 13.4 | 15.7 | 45.1 | 0 | | | | | |
| H3O+ | | | -76.867 | -76.716 | -76.731 | -96.4 | 21.5 | 23.9 | 48.4 | 0 | | | | | |
| O2 | | | -150.324 | -150.324 | -150.309 | -2.0 | 2.3 | 4.4 | 46.8 | 0 | | | | | |
| O4S-2 | | | -699.504 | -699.129 | -699.116 | -237.4 | 9.0 | 12.3 | 68.8 | 0 | | | | | |
| Oeeee | 63 | | -1153.129 | -1153.099 | -1153.488 | -20.8 | 276.5 | 291.7 | 159.2 | 0 | X | X | F | | Blue |
| oeeOe-R | 70 | | -1228.369 | -1228.332 | -1228.725 | -25.5 | 280.0 | 295.6 | 162.7 | 0 | X | I | F | | Blue |
| Oeeoe-R | 68 | | -1228.370 | -1228.334 | -1228.726 | -25.0 | 280.4 | 296.5 | 168.9 | 0 | X | I | F | | Blue |
| oeeOe-S | 71 | | -1228.372 | -1228.337 | -1228.726 | -24.2 | 279.8 | 296.1 | 173.6 | 0 | X | I | F | | Blue |
| Oeeoe-S | 69 | | -1228.364 | -1228.331 | -1228.725 | -22.3 | 280.4 | 296.3 | 165.2 | 0 | X | I | F | | Blue |
| oeeXe-R | 88 | | -1302.455 | -1302.408 | -1302.775 | -31.8 | 267.4 | 284.5 | 180.3 | 0 | X | X | F | | Blue |
| Oeexe-R | 84 | | -1302.461 | -1302.412 | -1302.785 | -32.7 | 268.1 | 284.3 | 168.4 | 0 | X | X | F | | Blue |
| oeeXe-S | 89 | | -1302.451 | -1302.406 | -1302.777 | -30.1 | 268.1 | 285.0 | 176.6 | 0 | X | X | F | | Blue |
| Oeexe-S | 85 | | -1302.459 | -1302.410 | -1302.784 | -32.6 | 268.5 | 284.6 | 167.9 | 0 | X | X | F | | Blue |
| Oeeye-R | 80 | | -1227.187 | -1227.154 | -1227.520 | -23.0 | 264.7 | 280.8 | 170.9 | 0 | X | X | F | | Blue |
| Oeeye-S | 81 | | -1227.187 | -1227.153 | -1227.518 | -23.2 | 264.5 | 280.6 | 171.4 | 0 | X | X | F | | Blue |
| Oeeyy | 102 | | -1301.225 | -1301.179 | -1301.525 | -31.0 | 252.6 | 269.2 | 173.7 | 0 | X | X | F | | Blue |
| oeOee | 67 | | -1228.372 | -1228.336 | -1228.730 | -24.6 | 279.4 | 294.9 | 159.9 | 0 | X | I | F | | Blue |
| Oeoee | 66 | | -1228.383 | -1228.344 | -1228.734 | -26.4 | 279.1 | 295.4 | 170.1 | 0 | X | I | F | | Blue |
| oOeee | 65 | | -1228.375 | -1228.331 | -1228.720 | -29.6 | 279.6 | 296.2 | 174.1 | 0 | X | X | F | | Blue |
| Ooeee | 64 | | -1228.373 | -1228.328 | -1228.717 | -30.0 | 279.4 | 296.1 | 175.1 | 0 | X | X | F | | Blue |
| ooeOe-R | 76 | | -1303.618 | -1303.563 | -1303.957 | -36.2 | 283.3 | 300.3 | 178.0 | 0 | X | X | X | I | Black |
| oOeoe-R | 74 | | -1303.620 | -1303.568 | -1303.961 | -34.4 | 283.1 | 300.1 | 179.6 | 0 | X | X | X | I | Black |
| Ooeoe-R | 72 | | -1303.609 | -1303.556 | -1303.948 | -35.1 | 282.6 | 300.0 | 180.4 | 0 | X | X | X | I | Black |
| ooeOe-S | 77 | | -1303.617 | -1303.564 | -1303.958 | -34.6 | 282.9 | 300.0 | 177.5 | 0 | X | X | X | I | Black |
| oOeoe-S | 75 | | -1303.615 | -1303.563 | -1303.959 | -34.3 | 283.7 | 300.6 | 176.2 | 0 | X | X | X | I | Black |
| Ooeoe-S | 73 | | -1303.602 | -1303.549 | -1303.944 | -35.3 | 283.4 | 300.4 | 175.9 | 0 | X | X | X | I | Black |
| ooeYe-5A-RR | 94 | | -1302.415 | -1302.364 | -1302.736 | -33.9 | 269.1 | 285.5 | 174.1 | 0 | X | X | F | | Blue |
| oOeye-5A-RR | 90 | | -1302.426 | -1302.377 | -1302.750 | -32.7 | 269.4 | 285.6 | 173.3 | 0 | X | X | F | | Blue |
| ooeYe-5A-RS | 95 | | -1302.417 | -1302.371 | -1302.748 | -30.2 | 270.0 | 285.9 | 165.9 | 0 | X | X | F | | Blue |
| oOeye-5A-RS | 91 | | -1302.425 | -1302.376 | -1302.749 | -32.8 | 269.5 | 285.7 | 173.4 | 0 | X | X | F | | Blue |
| ooeYe-5A-SR | 96 | | -1302.428 | -1302.379 | -1302.751 | -32.5 | 269.2 | 285.6 | 174.7 | 0 | X | X | F | | Blue |
| oOeye-5A-SR | 92 | | -1302.427 | -1302.378 | -1302.752 | -32.6 | 269.4 | 285.6 | 171.9 | 0 | X | X | F | | Blue |
| ooeYe-5A-SS | 97 | | -1302.419 | -1302.372 | -1302.745 | -31.9 | 269.2 | 285.5 | 171.9 | 0 | X | X | F | | Blue |
| oOeye-5A-SS | 93 | | -1302.428 | -1302.378 | -1302.753 | -33.1 | 269.3 | 285.0 | 165.4 | 0 | X | X | F | | Blue |
| oXeee-R | 86 | | -1302.443 | -1302.391 | -1302.762 | -34.4 | 268.2 | 285.0 | 175.3 | 0 | X | X | F | | Blue |
| Oxeee-R | 82 | | -1302.455 | -1302.401 | -1302.772 | -35.7 | 268.1 | 284.9 | 175.6 | 0 | X | X | F | | Blue |
| oXeee-S | 87 | | -1302.441 | -1302.390 | -1302.759 | -33.6 | 267.6 | 284.6 | 177.4 | 0 | X | X | F | | Blue |
| Oxeee-S | 83 | | -1302.460 | -1302.409 | -1302.777 | -33.6 | 267.8 | 284.7 | 181.0 | 0 | X | X | F | | Blue |
| Oyeee-R | 78 | | -1227.182 | -1227.140 | -1227.504 | -28.7 | 264.4 | 280.7 | 173.9 | 0 | X | X | F | | Blue |
| Oyeee-S | 79 | | -1227.182 | -1227.140 | -1227.504 | -28.4 | 264.0 | 280.4 | 174.7 | 0 | X | X | F | | Blue |
| Oyeye-RR | 98 | | -1301.228 | -1301.181 | -1301.524 | -31.7 | 252.1 | 269.0 | 179.8 | 0 | X | X | F | | Blue |
| Oyeye-RS | 99 | | -1301.239 | -1301.189 | -1301.536 | -33.3 | 252.6 | 269.0 | 172.9 | 0 | X | X | F | | Blue |
| Oyeye-SR | 100 | | -1301.234 | -1301.184 | -1301.529 | -33.0 | 252.5 | 269.3 | 176.9 | 0 | X | X | F | | Blue |
| Oyeye-SS | 101 | | -1301.232 | -1301.190 | -1301.536 | -28.5 | 252.7 | 269.2 | 173.8 | 0 | X | X | F | | Blue |
| oeeee | 1 | | -468.238 | -468.236 | -468.484 | -3.3 | 179.3 | 187.8 | 107.8 | 0 | F | F | F | | Red |
| oeeoe-R | 4 | | -543.483 | -543.473 | -543.726 | -8.1 | 183.1 | 192.0 | 111.5 | 0 | F | F | I | | Green |
| oeeoe-S | 5 | | -543.481 | -543.471 | -543.724 | -8.0 | 183.1 | 192.0 | 112.2 | 0 | F | F | I | | Green |
| oeeoo | 11 | | -618.726 | -618.707 | -618.963 | -13.8 | 186.5 | 196.1 | 117.0 | 0 | X | X | X | I | Black |
| oeexe-R | 18 | | -617.575 | -617.558 | -617.791 | -12.6 | 171.5 | 180.7 | 115.3 | 0 | X | F | F | | Red |
| oeexe-S | 19 | | -617.572 | -617.556 | -617.790 | -12.1 | 171.6 | 180.8 | 114.0 | 0 | X | F | F | | Red |
| oeeye-R | 14 | | -542.294 | -542.283 | -542.512 | -8.6 | 167.7 | 176.4 | 110.4 | 0 | X | X | I | | Gray |
| oeeye-S | 15 | | -542.294 | -542.283 | -542.511 | -8.5 | 167.5 | 176.4 | 111.7 | 0 | X | X | I | | Gray |
| oeeyo-5A-RR | 40 | | -617.534 | -617.521 | -617.758 | -10.0 | 173.0 | 181.6 | 108.9 | 0 | X | X | F | | Blue |
| oeeyo-5A-RS | 41 | | -617.537 | -617.519 | -617.756 | -13.2 | 172.7 | 181.4 | 111.3 | 0 | X | X | F | | Blue |
| oeeyo-5A-SR | 42 | | -617.532 | -617.515 | -617.752 | -12.4 | 172.6 | 181.4 | 111.6 | 0 | X | X | F | | Blue |
| oeeyo-5A-SS | 43 | | -617.530 | -617.515 | -617.751 | -11.3 | 172.8 | 181.5 | 110.7 | 0 | X | X | F | | Blue |
| oeeyo-R | 30 | | -617.534 | -617.520 | -617.753 | -11.0 | 171.5 | 180.6 | 114.2 | 0 | X | X | X | I | Black |
| oeeyo-S | 31 | | -617.533 | -617.518 | -617.752 | -11.6 | 171.5 | 180.7 | 114.5 | 0 | X | X | X | I | Black |
| oeeyy | 50 | | -616.342 | -616.327 | -616.534 | -11.4 | 155.5 | 164.8 | 116.3 | 0 | X | X | F | | Blue |
| oeoee | 3 | | -543.490 | -543.480 | -543.733 | -7.9 | 182.5 | 191.4 | 109.6 | 0 | F | F | I | | Green |
| oeoee-4D | 51 | | -467.003 | -467.000 | -467.228 | -3.7 | 165.2 | 172.8 | 100.5 | 0 | I | X | X | | Gray |
| oeooe-R | 9 | | -618.730 | -618.716 | -618.973 | -10.3 | 185.9 | 195.4 | 115.1 | 0 | X | X | X | I | Black |
| oeooe-S | 10 | | -618.730 | -618.711 | -618.967 | -13.6 | 185.7 | 195.3 | 116.1 | 0 | X | X | X | I | Black |
| oeoye-5A-RR | 36 | | -617.539 | -617.525 | -617.760 | -10.8 | 171.7 | 180.6 | 110.1 | 0 | X | X | F | | Blue |
| oeoye-5A-RS | 37 | | -617.539 | -617.528 | -617.764 | -8.9 | 171.9 | 180.7 | 109.3 | 0 | X | X | F | | Blue |
| oeoye-5A-SR | 38 | | -617.539 | -617.526 | -617.762 | -9.8 | 172.0 | 180.7 | 109.6 | 0 | X | X | F | | Blue |
| oeoye-5A-SS | 39 | | -617.538 | -617.523 | -617.759 | -11.3 | 171.9 | 180.6 | 110.3 | 0 | X | X | F | | Blue |
| oeoye-R | 28 | | -617.535 | -617.523 | -617.755 | -9.1 | 170.2 | 179.6 | 114.9 | 0 | X | X | X | I | Black |
| oeoye-S | 29 | | -617.538 | -617.522 | -617.754 | -11.4 | 170.0 | 179.6 | 115.7 | 0 | X | X | X | I | Black |
| ooeee | 2 | | -543.483 | -543.470 | -543.721 | -10.2 | 182.5 | 191.8 | 114.3 | 0 | X | F | I | | Green |
| ooeoe-R | 7 | | -618.726 | -618.705 | -618.963 | -14.9 | 186.8 | 196.3 | 116.3 | 0 | X | X | F | | Blue |
| ooeoe-S | 8 | | -618.724 | -618.704 | -618.960 | -14.5 | 186.4 | 196.0 | 117.6 | 0 | X | X | F | | Blue |
| ooeye-5A-RR | 32 | | -617.533 | -617.515 | -617.752 | -13.2 | 173.0 | 181.7 | 110.6 | 0 | X | X | F | | Blue |
| ooeye-5A-RS | 33 | | -617.532 | -617.514 | -617.751 | -12.9 | 172.9 | 181.5 | 111.1 | 0 | X | X | F | | Blue |
| ooeye-5A-SR | 34 | | -617.534 | -617.518 | -617.754 | -12.2 | 172.6 | 181.4 | 111.3 | 0 | X | X | F | | Blue |
| ooeye-5A-SS | 35 | | -617.536 | -617.519 | -617.756 | -12.6 | 172.7 | 181.4 | 111.0 | 0 | X | X | F | | Blue |
| ooeye-R | 26 | | -617.538 | -617.517 | -617.749 | -15.3 | 170.8 | 180.3 | 116.9 | 0 | X | X | X | I | Black |
| ooeye-S | 27 | | -617.538 | -617.517 | -617.748 | -15.3 | 170.6 | 180.2 | 118.5 | 0 | X | X | X | I | Black |
| oooee | 6 | | -618.734 | -618.716 | -618.971 | -13.7 | 185.5 | 195.2 | 116.5 | 0 | X | F | X | | Green |
| oooee-4D | 52 | | -542.248 | -542.234 | -542.465 | -10.4 | 168.4 | 176.8 | 106.8 | 0 | X | X | X | I | Black |
| oxeee-R | 16 | | -617.572 | -617.550 | -617.781 | -15.8 | 171.2 | 180.6 | 118.2 | 0 | X | F | F | | Red |
| oxeee-S | 17 | | -617.569 | -617.548 | -617.780 | -15.1 | 171.2 | 180.6 | 118.5 | 0 | X | F | F | | Red |
| oxoee-4D-R | 55 | | -616.335 | -616.314 | -616.525 | -15.2 | 156.9 | 165.5 | 110.3 | 0 | F | X | X | | Green |
| oxoee-4D-S | 56 | | -616.329 | -616.307 | -616.518 | -15.7 | 156.9 | 165.6 | 111.2 | 0 | F | X | X | | Green |
| oxoee-R | 44 | | -692.823 | -692.793 | -693.029 | -20.8 | 174.2 | 184.1 | 120.4 | 0 | F | F | X | | Green |
| oxoee-S | 45 | | -692.820 | -692.791 | -693.028 | -20.0 | 174.4 | 184.1 | 118.9 | 0 | F | F | X | | Green |
| oyeee-R | 12 | | -542.292 | -542.281 | -542.509 | -8.6 | 167.4 | 176.3 | 111.9 | 0 | X | X | I | | Gray |
| oyeee-S | 13 | | -542.290 | -542.280 | -542.508 | -8.1 | 167.7 | 176.5 | 111.1 | 0 | X | X | I | | Gray |
| oyeoe-RR | 22 | | -617.533 | -617.516 | -617.748 | -12.7 | 171.1 | 180.5 | 116.3 | 0 | X | X | X | I | Black |
| oyeoe-RS | 23 | | -617.534 | -617.516 | -617.748 | -13.3 | 171.1 | 180.4 | 117.0 | 0 | X | X | X | I | Black |
| oyeoe-SR | 24 | | -617.531 | -617.513 | -617.747 | -12.7 | 171.4 | 180.7 | 115.4 | 0 | X | X | X | I | Black |
| oyeoe-SS | 25 | | -617.532 | -617.513 | -617.746 | -13.5 | 171.3 | 180.5 | 114.9 | 0 | X | X | X | I | Black |
| oyeye-RR | 46 | | -616.346 | -616.328 | -616.534 | -13.3 | 155.4 | 164.8 | 118.9 | 0 | X | X | F | | Blue |
| oyeye-RS | 47 | | -616.347 | -616.328 | -616.536 | -14.2 | 155.7 | 164.9 | 116.0 | 0 | X | X | F | | Blue |
| oyeye-SR | 48 | | -616.343 | -616.326 | -616.534 | -12.5 | 155.8 | 165.0 | 116.0 | 0 | X | X | F | | Blue |
| oyeye-SS | 49 | | -616.344 | -616.325 | -616.532 | -13.7 | 155.3 | 164.7 | 117.6 | 0 | X | X | F | | Blue |
| oyoee-4D-R | 53 | | -541.057 | -541.045 | -541.252 | -9.1 | 153.3 | 161.3 | 105.3 | 0 | X | X | X | I | Black |
| oyoee-4D-S | 54 | | -541.055 | -541.044 | -541.251 | -9.0 | 153.4 | 161.3 | 104.6 | 0 | X | X | X | I | Black |
| oyoee-R | 20 | | -617.543 | -617.525 | -617.757 | -13.4 | 170.7 | 179.9 | 114.0 | 0 | X | X | X | I | Black |

| | | | | | | | | | | | | | | | |
|---|---|---|---|---|---|---|---|---|---|---|---|---|---|---|---|
| oyoee-S | **21** | | -617.542 | -617.525 | -617.757 | -12.7 | 170.6 | 179.9 | 114.4 | 0 | X | X | X | I | **Black** |
| seeee | **57** | | -1092.129 | -1092.116 | -1092.373 | -9.9 | 188.9 | 199.9 | 130.2 | 0 | X | X | F | | **Blue** |
| seeoe-R | **59** | | -1167.367 | -1167.344 | -1167.606 | -16.0 | 192.3 | 203.9 | 133.2 | 0 | X | X | F | | **Blue** |
| seeoe-S | **60** | | -1167.370 | -1167.349 | -1167.608 | -15.2 | 191.8 | 203.6 | 137.3 | 0 | X | X | F | | **Blue** |
| soeee | **58** | | -1167.372 | -1167.348 | -1167.608 | -16.9 | 192.0 | 203.8 | 137.5 | 0 | X | X | F | | **Blue** |
| soeoe-R | **61** | | -1242.606 | -1242.575 | -1242.840 | -21.6 | 195.6 | 207.8 | 139.5 | 0 | X | X | F | | **Blue** |
| soeoe-S | **62** | | -1242.607 | -1242.576 | -1242.840 | -21.6 | 195.4 | 207.8 | 141.4 | 0 | X | X | F | | **Blue** |
| | | | | | | | | | | X | 93 | 84 | 30 | | |
| | | | | | | | | | | F | 8 | 12 | 64 | | |
| | | | | | | | | | | I | 1 | 6 | 8 | 24 | |
| | | | | | | | | | | **Red** | | | | | **5** |
| | | | | | | | | | | **Green** | | | | | **9** |
| | | | | | | | | | | **Blue** | | | | | **59** |
| | | | | | | | | | | **Gray** | | | | | **5** |
| | | | | | | | | | | **Black** | | | | | **24** |
| | | | | | | | | | | Total | 102 | 102 | 102 | 24 | 102 |

| Hess's law | | | | | | | | | | | | |
|---|---|---|---|---|---|---|---|---|---|---|---|---|
| **Reactions** | **$\Delta G^{*}_{soln}$** | **$\Delta G°_{gas}$** | | | | **$\Delta\Delta G_{solv}+\Delta\Delta G$ o->*** | | | | **STEP** | **RXN** | **POS** |
| | **kcal/mol** | **kcal/mol** | | | | **kcal/mol** | | | | | | |
| C6H10O7 -> H2O | 429778.3 | 429754.9 | | | | 23.4 | | | | | | |
| 2H3O+ + SO4-2 -> 3H2O | 391473.2 | 391061.7 | | | | 411.5 | | | | | | |
| 1/2O2 -> 0O2 | 47165.0 | 47164.9 | | | | 0.1 | | | | | | |
| 0H2 -> H2 | -737.6 | -741.1 | | | | 3.5 | | | | | | |
| 0HO2 -> H2O | -47980.5 | -47976.2 | | | | -4.3 | | | | | | |
| oeeee + C6H10O7 -> H2O + Oeeee | 3.0 | -2.9 | | | | 5.9 | | | | 1 | g | 3 |
| ooeee + C6H10O7 -> H2O + Ooeee | 3.4 | -0.1 | | | | 3.6 | | | | 2 | g | 3 |
| ooeee + C6H10O7 -> H2O + oOeee | 2.2 | -1.7 | | | | 3.9 | | | | 2 | g | 7 |
| oeoee + C6H10O7 -> H2O + Oeoee | 1.2 | -3.7 | | | | 4.9 | | | | 2 | g | 3 |
| oeoee + C6H10O7 -> H2O + oeOee | 8.3 | 1.6 | | | | 6.7 | | | | 2 | g | 8 |
| oeeoe-R + C6H10O7 -> H2O + Oeeoe-R | 4.9 | -1.6 | | | | 6.5 | | | | 2 | g | 3 |
| oeeoe-S + C6H10O7 -> H2O + Oeeoe-S | 7.5 | -1.6 | | | | 9.1 | | | | 2 | g | 3 |
| oeeoe-R + C6H10O7 -> H2O + oeeOe-R | 5.7 | -0.3 | | | | 6.0 | | | | 2 | g | 9 |
| oeeoe-S + C6H10O7 -> H2O + oeeOe-S | 2.3 | -4.9 | | | | 7.2 | | | | 2 | g | 9 |
| oyeee-R + C6H10O7 -> H2O + Oyeee-R | 3.0 | -0.3 | | | | 3.3 | | | | 3 | g | 3 |
| oyeee-S + C6H10O7 -> H2O + Oyeee-S | 1.9 | -1.2 | | | | 3.1 | | | | 3 | g | 3 |
| oeeye-R + C6H10O7 -> H2O + Oeeye-R | 1.2 | -7.7 | | | | 8.9 | | | | 3 | g | 3 |
| oeeye-S + C6H10O7 -> H2O + Oeeye-S | 1.5 | -7.2 | | | | 8.7 | | | | 3 | g | 3 |
| ooeoe-R + C6H10O7 -> H2O + Ooeoe-R | 7.9 | 4.7 | | | | 3.2 | | | | 3 | g | 3 |
| ooeoe-S + C6H10O7 -> H2O + Ooeoe-S | 10.7 | 8.1 | | | | 2.6 | | | | 3 | g | 3 |
| ooeoe-R + C6H10O7 -> H2O + oOeoe-R | 0.8 | -3.1 | | | | 3.9 | | | | 3 | g | 7 |
| ooeoe-S + C6H10O7 -> H2O + oOeoe-S | 2.7 | -0.9 | | | | 3.6 | | | | 3 | g | 7 |
| ooeoe-R + C6H10O7 -> H2O + ooeOe-R | 2.1 | 0.0 | | | | 2.1 | | | | 3 | g | 9 |
| ooeoe-S + C6H10O7 -> H2O + ooeOe-S | 1.8 | -1.5 | | | | 3.3 | | | | 3 | g | 9 |
| oxeee-R + C6H10O7 -> H2O + Oxeee-R | 7.4 | 3.9 | | | | 3.4 | | | | 4 | g | 3 |
| oxeee-S + C6H10O7 -> H2O + Oxeee-S | 3.1 | -1.7 | | | | 4.9 | | | | 4 | g | 3 |
| oeexe-R + C6H10O7 -> H2O + Oeexe-R | 5.8 | 2.5 | | | | 3.3 | | | | 4 | g | 3 |
| oeexe-S + C6H10O7 -> H2O + Oeexe-S | 4.9 | 2.1 | | | | 2.8 | | | | 4 | g | 3 |
| oxeee-R + C6H10O7 -> H2O + oXeee-R | 15.0 | 10.3 | | | | 4.7 | | | | 4 | g | 7 |
| oxeee-S + C6H10O7 -> H2O + oXeee-S | 15.1 | 10.2 | | | | 4.9 | | | | 4 | g | 7 |
| oeexe-R + C6H10O7 -> H2O + oeeXe-R | 9.3 | 5.2 | | | | 4.1 | | | | 4 | g | 9 |
| oeexe-S + C6H10O7 -> H2O + oeeXe-S | 10.0 | 4.6 | | | | 5.3 | | | | 4 | g | 9 |
| ooeye-5A-RR + C6H10O7 -> H2O + oOeye-5A-RR | 1.5 | -2.5 | | | | 3.9 | | | | 5 | g | 7 |
| ooeye-5A-RS + C6H10O7 -> H2O + oOeye-5A-RS | 1.3 | -2.2 | | | | 3.5 | | | | 5 | g | 7 |
| ooeye-5A-SR + C6H10O7 -> H2O + oOeye-5A-SR | 1.3 | -1.6 | | | | 2.9 | | | | 5 | g | 7 |
| ooeye-5A-SS + C6H10O7 -> H2O + oOeye-5A-SS | 2.6 | -0.3 | | | | 2.9 | | | | 5 | g | 7 |
| ooeye-5A-RS + C6H10O7 -> H2O + ooeYe-5A-RR | 7.8 | 5.4 | | | | 2.4 | | | | 5 | g | 9 |
| ooeye-5A-RR + C6H10O7 -> H2O + ooeYe-5A-RS | 7.5 | 1.1 | | | | 6.4 | | | | 5 | g | 9 |
| ooeye-5A-SS + C6H10O7 -> H2O + ooeYe-5A-SR | 2.3 | -1.1 | | | | 3.5 | | | | 5 | g | 9 |
| ooeye-5A-SR + C6H10O7 -> H2O + ooeYe-5A-SS | 6.3 | 2.7 | | | | 3.6 | | | | 5 | g | 9 |
| oyeye-RR + C6H10O7 -> H2O + Oyeye-RR | 8.2 | 3.3 | | | | 5.0 | | | | 5 | g | 3 |
| oyeye-RS + C6H10O7 -> H2O + Oyeye-RS | 2.3 | -2.0 | | | | 4.3 | | | | 5 | g | 3 |
| oyeye-SR + C6H10O7 -> H2O + Oyeye-SR | 3.0 | 0.1 | | | | 2.9 | | | | 5 | g | 3 |
| oyeye-SS + C6H10O7 -> H2O + Oyeye-SS | 4.3 | -4.3 | | | | 8.6 | | | | 5 | g | 3 |
| oeeyy + C6H10O7 -> H2O + Oeeyy | 7.7 | 4.0 | | | | 3.7 | | | | 5 | g | 3 |
| oeeee + 2H3O+ + SO4-2 -> 3H2O + seeee | -24.0 | -428.9 | | | | 404.9 | | | | 1 | s | 3 |
| ooeee + 2H3O+ + SO4-2 -> 3H2O + soeee | -23.0 | -427.7 | | | | 404.7 | | | | 2 | s | 3 |
| oeeoe-R + 2H3O+ + SO4-2 -> 3H2O + seeoe-R | -19.9 | -423.5 | | | | 403.6 | | | | 2 | s | 3 |
| oeeoe-S + 2H3O+ + SO4-2 -> 3H2O + seeoe-S | -23.4 | -427.7 | | | | 404.3 | | | | 2 | s | 3 |
| ooeoe-R + 2H3O+ + SO4-2 -> 3H2O + soeoe-R | -17.5 | -422.3 | | | | 404.8 | | | | 3 | s | 3 |
| ooeoe-S + 2H3O+ + SO4-2 -> 3H2O + soeoe-S | -19.2 | -423.6 | | | | 404.4 | | | | 3 | s | 3 |
| oeeee + 1/2O2 -> ooeee | -52.1 | -45.3 | | | | -6.8 | | | | 1 | o1 | 7 |
| oeeee + 1/2O2 -> oeoee | -56.0 | -51.5 | | | | -4.5 | | | | 1 | o1 | 8 |
| oeeee + 1/2O2 -> oeeoe-R | -51.9 | -47.2 | | | | -4.8 | | | | 1 | o1 | 9 |
| oeeee + 1/2O2 -> oeeoe-S | -50.4 | -45.7 | | | | -4.7 | | | | 1 | o1 | 9 |
| Oeeee + 1/2O2 -> Ooeee | -51.7 | -42.5 | | | | -9.2 | | | | 2 | o1 | 7 |
| Oeeee + 1/2O2 -> Oeoee | -57.8 | -52.3 | | | | -5.5 | | | | 2 | o1 | 8 |
| Oeeee + 1/2O2 -> Oeeoe-R | -50.0 | -45.8 | | | | -4.2 | | | | 2 | o1 | 9 |
| Oeeee + 1/2O2 -> Oeeoe-S | -45.9 | -44.4 | | | | -1.4 | | | | 2 | o1 | 9 |
| ooeee + 1/2O2 -> oooee | -55.7 | -52.3 | | | | -3.5 | | | | 2 | o1 | 8 |
| oeoee + 1/2O2 -> oooee | -51.8 | -46.1 | | | | -5.7 | | | | 2 | o1 | 7 |
| ooeee + 1/2O2 -> ooeoe-R | -50.5 | -45.9 | | | | -4.7 | | | | 2 | o1 | 9 |
| ooeee + 1/2O2 -> ooeoe-S | -49.3 | -45.0 | | | | -4.3 | | | | 2 | o1 | 9 |
| oeeoe-R + 1/2O2 -> ooeoe-R | -50.7 | -44.0 | | | | -6.7 | | | | 2 | o1 | 7 |
| oeeoe-S + 1/2O2 -> ooeoe-S | -51.0 | -44.6 | | | | -6.4 | | | | 2 | o1 | 7 |
| oeoee + 1/2O2 -> oeooe-R | -48.8 | -46.5 | | | | -2.3 | | | | 2 | o1 | 9 |
| oeoee + 1/2O2 -> oeooe-S | -48.9 | -43.3 | | | | -5.7 | | | | 2 | o1 | 9 |
| oeeoe-S + 1/2O2 -> oeooe-R | -54.5 | -52.3 | | | | -2.2 | | | | 2 | o1 | 8 |
| oeeoe-R + 1/2O2 -> oeooe-S | -53.0 | -47.6 | | | | -5.4 | | | | 2 | o1 | 8 |
| oeeoe-R + 1/2O2 -> oeeoo | -50.5 | -44.9 | | | | -5.6 | | | | 2 | o1 | 10 |
| oeeoe-S + 1/2O2 -> oeeoo | -52.0 | -46.3 | | | | -5.7 | | | | 2 | o1 | 10 |
| Ooeee + 1/2O2 -> Ooeoe-R | -46.1 | -41.1 | | | | -5.0 | | | | 3 | o1 | 9 |
| Ooeee + 1/2O2 -> Ooeoe-S | -42.0 | -36.8 | | | | -5.2 | | | | 3 | o1 | 9 |
| Oeeoe-R + 1/2O2 -> Ooeoe-R | -47.8 | -37.8 | | | | -10.0 | | | | 3 | o1 | 7 |
| Oeeoe-S + 1/2O2 -> Ooeoe-S | -47.8 | -34.9 | | | | -12.9 | | | | 3 | o1 | 7 |
| oOeee + 1/2O2 -> oOeoe-R | -52.0 | -47.3 | | | | -4.7 | | | | 3 | o1 | 9 |
| oOeee + 1/2O2 -> oOeoe-S | -48.8 | -44.2 | | | | -4.6 | | | | 3 | o1 | 9 |
| oeeOe-R + 1/2O2 -> ooeOe-R | -54.4 | -43.7 | | | | -10.6 | | | | 3 | o1 | 7 |
| oeeOe-S + 1/2O2 -> ooeOe-S | -51.5 | -41.2 | | | | -10.4 | | | | 3 | o1 | 7 |
| oeoee-4D + 1/2O2 -> oooee-4D | -51.9 | -45.2 | | | | -6.7 | | | | 3 | o1 | 7 |
| oyeee-R + 1/2O2 -> oyoee-R | -55.8 | -51.0 | | | | -4.7 | | | | 3 | o1 | 8 |
| oyeee-S + 1/2O2 -> oyoee-S | -56.4 | -51.8 | | | | -4.5 | | | | 3 | o1 | 8 |
| oyeee-R + 1/2O2 -> oyeoe-RR | -49.5 | -45.5 | | | | -4.0 | | | | 3 | o1 | 9 |
| oyeee-R + 1/2O2 -> oyeoe-RS | -50.3 | -45.6 | | | | -4.7 | | | | 3 | o1 | 9 |
| oyeee-S + 1/2O2 -> oyeoe-SR | -49.4 | -44.9 | | | | -4.5 | | | | 3 | o1 | 9 |
| oyeee-S + 1/2O2 -> oyeoe-SS | -50.1 | -44.8 | | | | -5.3 | | | | 3 | o1 | 9 |
| oeeye-R + 1/2O2 -> ooeye-R | -51.6 | -44.9 | | | | -6.7 | | | | 3 | o1 | 7 |
| oeeye-S + 1/2O2 -> ooeye-S | -51.8 | -45.0 | | | | -6.7 | | | | 3 | o1 | 7 |
| oeeye-S + 1/2O2 -> oeoye-R | -49.5 | -48.9 | | | | -0.6 | | | | 3 | o1 | 8 |
| oeeye-R + 1/2O2 -> oeoye-S | -51.0 | -48.2 | | | | -2.8 | | | | 3 | o1 | 8 |
| oeeye-R + 1/2O2 -> oeeyo-R | -49.0 | -46.7 | | | | -2.4 | | | | 3 | o1 | 10 |
| oeeye-S + 1/2O2 -> oeeyo-S | -48.7 | -45.7 | | | | -3.0 | | | | 3 | o1 | 10 |
| oxeee-S + 1/2O2 -> oxoee-S | -55.3 | -50.5 | | | | -4.8 | | | | 4 | o1 | 8 |
| oxeee-R + 1/2O2 -> oxoee-R | -55.9 | -51.0 | | | | -4.9 | | | | 4 | o1 | 8 |
| oyeee-R + 1/2O2 -> oxeee-R | -73.8 | -66.7 | | | | -7.1 | | | | 3 | o2 | 7 |
| oyeee-S + 1/2O2 -> oxeee-S | -73.6 | -66.8 | | | | -6.9 | | | | 3 | o2 | 7 |
| oeeye-R + 1/2O2 -> oeexe-R | -74.3 | -70.4 | | | | -3.9 | | | | 3 | o2 | 9 |

| | | | | | | | | | | | | |
|---|---|---|---|---|---|---|---|---|---|---|---|---|
| oeeye-S + 1/2O2 -> oeexe-S | -72.8 | -69.3 | | | | -3.5 | | | | 3 | o2 | 9 |
| Oyeee-R + 1/2O2 -> Oxeee-R | -69.4 | -62.5 | | | | -6.9 | | | | 4 | o2 | 7 |
| Oyeee-S + 1/2O2 -> Oxeee-S | -72.4 | -67.3 | | | | -5.1 | | | | 4 | o2 | 7 |
| Oeeye-R + 1/2O2 -> Oeexe-R | -69.8 | -60.2 | | | | -9.6 | | | | 4 | o2 | 9 |
| Oeeye-S + 1/2O2 -> Oeexe-S | -69.4 | -60.0 | | | | -9.4 | | | | 4 | o2 | 9 |
| oyoee-R + 1/2O2 -> oxoee-R | -74.0 | -66.7 | | | | -7.3 | | | | 4 | o2 | 7 |
| oyoee-S + 1/2O2 -> oxoee-S | -72.6 | -65.4 | | | | -7.2 | | | | 4 | o2 | 7 |
| oyoee-4D-R + 1/2O2 -> oxoee-4D-R | -72.9 | -67.0 | | | | -6.0 | | | | 5 | o2 | 7 |
| oyoee-4D-S + 1/2O2 -> oxoee-4D-S | -70.2 | -63.5 | | | | -6.7 | | | | 5 | o2 | 7 |
| ooeee -> oyeee-R + H2 | 10.1 | 5.0 | | | | 5.1 | | | | 2 | o3 | 7 |
| ooeee -> oyeee-S + H2 | 11.4 | 5.8 | | | | 5.6 | | | | 2 | o3 | 7 |
| oeeoe-R -> oeeye-R + H2 | 8.5 | 5.4 | | | | 3.1 | | | | 2 | o3 | 9 |
| oeeoe-S -> oeeye-S + H2 | 7.2 | 4.1 | | | | 3.1 | | | | 2 | o3 | 9 |
| Ooeee -> Oyeee-R + H2 | 9.7 | 4.8 | | | | 4.8 | | | | 3 | o3 | 7 |
| Ooeee -> Oyeee-S + H2 | 9.9 | 4.8 | | | | 5.1 | | | | 3 | o3 | 7 |
| Oeeoe-R -> Oeeye-R + H2 | 4.8 | -0.7 | | | | 5.5 | | | | 3 | o3 | 9 |
| Oeeoe-S -> Oeeye-S + H2 | 1.2 | -1.4 | | | | 2.6 | | | | 3 | o3 | 9 |
| oooee -> oyoee-R + H2 | 10.1 | 6.3 | | | | 3.8 | | | | 3 | o3 | 7 |
| oooee -> oyoee-S + H2 | 10.7 | 6.2 | | | | 4.5 | | | | 3 | o3 | 7 |
| ooeoe-R -> oyeoe-RR + H2 | 11.2 | 5.4 | | | | 5.8 | | | | 3 | o3 | 7 |
| ooeoe-R -> oyeoe-RS + H2 | 10.4 | 5.2 | | | | 5.1 | | | | 3 | o3 | 7 |
| ooeoe-S -> oyeoe-SR + H2 | 11.2 | 5.9 | | | | 5.3 | | | | 3 | o3 | 7 |
| ooeoe-S -> oyeoe-SS + H2 | 10.5 | 6.0 | | | | 4.5 | | | | 3 | o3 | 7 |
| ooeoe-R -> ooeye-R + H2 | 7.7 | 4.5 | | | | 3.2 | | | | 3 | o3 | 9 |
| ooeoe-S -> ooeye-S + H2 | 6.4 | 3.7 | | | | 2.7 | | | | 3 | o3 | 9 |
| oeooe-R -> oeoye-R + H2 | 12.2 | 7.5 | | | | 4.7 | | | | 3 | o3 | 9 |
| oeooe-S -> oeoye-S + H2 | 10.5 | 4.8 | | | | 5.7 | | | | 3 | o3 | 9 |
| oeeoo -> oeeyo-R + H2 | 9.9 | 3.6 | | | | 6.3 | | | | 3 | o3 | 9 |
| oeeoo -> oeeyo-S + H2 | 10.5 | 4.8 | | | | 5.7 | | | | 3 | o3 | 9 |
| oooee-4D -> oyoee-4D-R + H2 | 9.9 | 5.1 | | | | 4.8 | | | | 4 | o3 | 7 |
| oooee-4D -> oyoee-4D-S + H2 | 11.0 | 6.0 | | | | 5.0 | | | | 4 | o3 | 7 |
| oyeoe-RR -> oyeye-RR + H2 | 7.2 | 4.3 | | | | 2.9 | | | | 4 | o3 | 9 |
| oyeoe-RS -> oyeye-RS + H2 | 7.1 | 4.4 | | | | 2.7 | | | | 4 | o3 | 9 |
| oyeoe-SR -> oyeye-SR + H2 | 7.6 | 3.8 | | | | 3.8 | | | | 4 | o3 | 9 |
| oyeoe-SS -> oyeye-SS + H2 | 8.0 | 4.6 | | | | 3.3 | | | | 4 | o3 | 9 |
| ooeye-R -> oyeye-RR + H2 | 10.7 | 5.2 | | | | 5.5 | | | | 4 | o3 | 7 |
| ooeye-R -> oyeye-SR + H2 | 12.4 | 6.1 | | | | 6.3 | | | | 4 | o3 | 7 |
| ooeye-S -> oyeye-RS + H2 | 9.8 | 5.1 | | | | 4.7 | | | | 4 | o3 | 7 |
| ooeye-S -> oyeye-SS + H2 | 12.1 | 7.0 | | | | 5.1 | | | | 4 | o3 | 7 |
| oeeyo-R -> oeeyy + H2 | 10.6 | 7.4 | | | | 3.2 | | | | 4 | o3 | 10 |
| oeeyo-S -> oeeyy + H2 | 10.0 | 6.3 | | | | 3.7 | | | | 4 | o3 | 10 |
| oeoee -> oeoee-4D + H2O | 15.6 | 15.6 | | | | -0.1 | | | | 2 | d | |
| oooee -> oooee-4D + H2O | 15.5 | 16.6 | | | | -1.1 | | | | 3 | d | |
| oyoee-R -> oyoee-4D-R + H2O | 15.3 | 15.4 | | | | -0.1 | | | | 4 | d | |
| oyoee-S -> oyoee-4D-S + H2O | 15.8 | 16.3 | | | | -0.6 | | | | 4 | d | |
| oxoee-R -> oxoee-4D-R + H2O | 16.3 | 15.1 | | | | 1.2 | | | | 5 | d | |
| oxoee-S -> oxoee-4D-S + H2O | 18.1 | 18.2 | | | | -0.1 | | | | 5 | d | |
| ooeye-R -> ooeye-5A-RR | 3.3 | 1.2 | | | | 2.1 | | | | 4 | a | |
| ooeye-R -> ooeye-5A-RS | 4.2 | 1.8 | | | | 2.4 | | | | 4 | a | |
| ooeye-S -> ooeye-5A-SR | 2.8 | -0.4 | | | | 3.1 | | | | 4 | a | |
| ooeye-S -> ooeye-5A-SS | 1.2 | -1.5 | | | | 2.7 | | | | 4 | a | |
| oeoye-R -> oeoye-5A-RR | -2.7 | -1.0 | | | | -1.6 | | | | 4 | a | |
| oeoye-R -> oeoye-5A-RS | -2.6 | -2.9 | | | | 0.2 | | | | 4 | a | |
| oeoye-S -> oeoye-5A-SR | -0.7 | -2.3 | | | | 1.6 | | | | 4 | a | |
| oeoye-S -> oeoye-5A-SS | -0.5 | -0.6 | | | | 0.1 | | | | 4 | a | |
| oeeyo-R -> oeeyo-5A-RR | 0.5 | -0.5 | | | | 1.0 | | | | 4 | a | |
| oeeyo-R -> oeeyo-5A-RS | -1.9 | 0.3 | | | | -2.2 | | | | 4 | a | |
| oeeyo-S -> oeeyo-5A-SR | 0.8 | 1.7 | | | | -0.9 | | | | 4 | a | |
| oeeyo-S -> oeeyo-5A-SS | 2.3 | 2.0 | | | | 0.3 | | | | 4 | a | |

| Step of reaction | Average | | SD | | |
|---|---|---|---|---|---|
| | $\Delta G^*_{soln}$ | $\Delta G°_{gas}$ | $\Delta G^*_{soln}$ | $\Delta G°_{gas}$ | |
| | kcal/mol | kcal/mol | kcal/mol | kcal/mol | |
| Step 1 | -38.6 | -103.6 | 23.4 | 160.4 | 1 |
| Step 2 | -25.0 | -62.5 | 28.2 | 121.2 | 2 |
| Step 3 | -22.0 | -36.6 | 31.1 | 80.9 | 3 |
| Step 4 | -7.5 | -8.4 | 30.0 | 26.0 | 4 |
| Step 5 | -3.1 | -5.6 | 26.2 | 23.3 | 5 |

| Type of reaction | Average | | SD | | |
|---|---|---|---|---|---|
| | $\Delta G^*_{soln}$ | $\Delta G°_{gas}$ | $\Delta G^*_{soln}$ | $\Delta G°_{gas}$ | |
| | kcal/mol | kcal/mol | kcal/mol | kcal/mol | |
| g | 5.0 | 0.4 | 3.7 | 4.1 | g |
| s | -21.2 | -425.6 | 2.7 | 2.8 | s |
| o1 | -51.1 | -45.8 | 3.1 | 3.9 | o1 |
| o2 | -72.1 | -65.5 | 1.9 | 3.3 | o2 |
| o3 | 9.4 | 4.9 | 2.3 | 1.9 | o3 |
| d | 16.1 | 16.2 | 1.0 | 1.1 | d |
| a | 0.6 | -0.2 | 2.3 | 1.6 | a |

| Position of reaction | Average | | SD | | |
|---|---|---|---|---|---|
| | $\Delta G^*_{soln}$ | $\Delta G°_{gas}$ | $\Delta G^*_{soln}$ | $\Delta G°_{gas}$ | |
| | kcal/mol | kcal/mol | kcal/mol | kcal/mol | |
| 3 | -1.3 | -98.4 | 11.4 | 182.8 | 3 |
| 7 | -21.8 | -20.9 | 35.2 | 29.3 | 7 |
| 8 | -49.4 | -46.3 | 18.3 | 15.2 | 8 |
| 9 | -20.6 | -20.7 | 31.1 | 26.6 | 9 |
| 10 | -29.9 | -28.3 | 31.2 | 27.2 | 10 |

| Type and position of reaction | Average | | SD | |
|---|---|---|---|---|
| | $\Delta G^*_{soln}$ | $\Delta G°_{gas}$ | $\Delta G^*_{soln}$ | $\Delta G°_{gas}$ |
| | kcal/mol | kcal/mol | kcal/mol | kcal/mol |
| g-3 | 4.7 | -0.3 | 2.8 | 4.0 |
| g-7 | 4.7 | 0.9 | 5.9 | 5.4 |
| g-8 | 8.3 | 1.6 | #DIV/0! | #DIV/0! |
| g-9 | 5.5 | 1.1 | 3.2 | 3.4 |
| s-3 | -21.2 | -425.6 | 2.7 | 2.8 |
| o1-7 | -51.2 | -42.9 | 1.8 | 0.6 |
| o1-8 | -54.6 | -50.7 | 2.5 | 1.7 |
| o1-9 | -49.0 | -44.6 | 2.5 | 2.6 |
| o1-10 | -50.0 | -45.9 | 1.5 | 0.8 |
| o2-7 | -72.4 | -65.7 | 1.7 | 1.8 |
| o2-9 | -71.6 | -65.0 | 2.4 | 5.6 |
| o3-7 | 10.7 | 5.6 | 0.8 | 0.6 |
| o3-9 | 7.8 | 3.8 | 2.7 | 2.3 |
| o3-10 | 10.3 | 6.8 | 0.4 | 0.8 |
| 4D | 16.1 | 16.2 | 1.0 | 1.1 |
| 5A | 0.6 | -0.2 | 2.3 | 1.6 |

Compounds and reactions

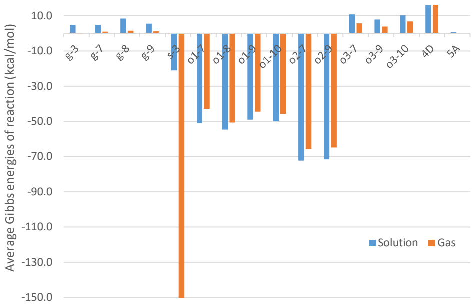

| name | step | energy (kcal/mol) | | reactions |
|---|---|---|---|---|
| | | aqueous phase | gas phase | |
| oeeee -> oeeee | 0 | 0.0 | 0.0 | |
| oeeee + 1/2O2 -> ooeee | 1 | -52.1 | -45.3 | o1 |
| oeeee + 1/2O2 -> oeoee | 1 | -56.0 | -51.5 | |
| oeeee + 1/2O2 -> oeeoe-R | 1 | -51.9 | -47.2 | |
| oeeee + 1/2O2 -> oeeoe-S | 1 | -50.4 | -45.7 | |
| oeeee + C6H10O7 -> H2O + Oeeee | 1 | 3.0 | -2.9 | g |
| oeeee + 2H3O+ + SO4-2 -> 3H2O + seeee | 1 | -24.0 | -428.9 | s |
| oeeee + O2 -> oooee | 2 | -107.9 | -97.6 | o1 x 2 |
| oeeee + O2 -> ooeoe-R | 2 | -102.7 | -91.2 | |
| oeeee + O2 -> ooeoe-S | 2 | -101.4 | -90.3 | |
| oeeee + O2 -> oeooe-R | 2 | -104.9 | -98.0 | |
| oeeee + O2 -> oeooe-S | 2 | -105.0 | -94.8 | |
| oeeee + O2 -> oeeoo | 2 | -102.4 | -92.0 | |
| oeeee + 1/2O2 -> oeoee-4D + H2O | 2 | -40.5 | -35.8 | o1, 4D |
| oeeee + 1/2O2 -> oyeee-R + H2 | 2 | -42.0 | -40.3 | o1, o3 |
| oeeee + 1/2O2 -> oyeee-S + H2 | 2 | -40.7 | -39.5 | |
| oeeee + 1/2O2 -> oeeye-R + H2 | 2 | -43.4 | -41.7 | |
| oeeee + 1/2O2 -> oeeye-S + H2 | 2 | -43.2 | -41.6 | |
| oeeee + C6H10O7 + 1/2O2 -> H2O + Ooeee | 2 | -48.7 | -45.4 | o1, g |
| oeeee + C6H10O7 + 1/2O2 -> H2O + oOeee | 2 | -49.9 | -47.0 | |
| oeeee + C6H10O7 + 1/2O2 -> H2O + Oeoee | 2 | -54.8 | -55.2 | |
| oeeee + C6H10O7 + 1/2O2 -> H2O + oeOee | 2 | -47.7 | -49.9 | |
| oeeee + C6H10O7 + 1/2O2 -> H2O + Oeeoe-R | 2 | -47.0 | -48.7 | |
| oeeee + C6H10O7 + 1/2O2 -> H2O + Oeeoe-S | 2 | -42.9 | -47.3 | |
| oeeee + C6H10O7 + 1/2O2 -> H2O + oeeOe-R | 2 | -46.2 | -47.4 | |
| oeeee + C6H10O7 + 1/2O2 -> H2O + oeeOe-S | 2 | -48.1 | -50.7 | |
| oeeee + 2H3O+ + SO4-2 + 1/2O2 -> 3H2O + soeee | 2 | -75.1 | -473.0 | o1, s |
| oeeee + 2H3O+ + SO4-2 + 1/2O2 -> 3H2O + seeoe-R | 2 | -71.8 | -470.6 | |
| oeeee + 2H3O+ + SO4-2 + 1/2O2 -> 3H2O + seeoe-S | 2 | -73.8 | -473.4 | |
| oeeee + O2 -> H2 + oxeee-R | 3 | -115.8 | -107.0 | o1, o3, o2 |
| oeeee + O2 -> H2 + oxeee-S | 3 | -114.4 | -106.3 | |
| oeeee + O2 -> H2 + oeexe-R | 3 | -117.7 | -112.1 | |
| oeeee + O2 -> H2 + oeexe-S | 3 | -116.0 | -110.9 | |
| oeeee + O2 -> H2 + oyoee-R | 3 | -97.8 | -91.3 | o1 x 2, o3 |
| oeeee + O2 -> H2 + oyoee-S | 3 | -97.1 | -91.3 | |
| oeeee + O2 -> H2 + oyeoe-RR | 3 | -91.5 | -85.8 | |
| oeeee + O2 -> H2 + oyeoe-RS | 3 | -92.3 | -85.9 | |
| oeeee + O2 -> H2 + oyeoe-SR | 3 | -90.1 | -84.4 | |
| oeeee + O2 -> H2 + oyeoe-SS | 3 | -90.9 | -84.3 | |
| oeeee + O2 -> H2 + ooeye-R | 3 | -95.0 | -86.6 | |
| oeeee + O2 -> H2 + ooeye-S | 3 | -95.0 | -86.6 | |
| oeeee + O2 -> H2 + oeoye-R | 3 | -92.7 | -90.5 | |
| oeeee + O2 -> H2 + oeoye-S | 3 | -94.4 | -90.0 | |
| oeeee + O2 -> H2 + oeeyo-R | 3 | -92.4 | -88.4 | |
| oeeee + O2 -> H2 + oeeyo-S | 3 | -91.9 | -87.3 | |
| oeeee + O2 -> H2O + oooee-4D | 3 | -92.3 | -81.0 | o1 x 2, 4D |
| oeeee + C6H10O7 + O2 -> H2O + Ooeoe-R | 3 | -94.8 | -86.5 | o1 x 2, g |
| oeeee + C6H10O7 + O2 -> H2O + Ooeoe-S | 3 | -90.7 | -82.2 | |
| oeeee + C6H10O7 + O2 -> H2O + oOeoe-R | 3 | -101.9 | -94.3 | |
| oeeee + C6H10O7 + O2 -> H2O + oOeoe-S | 3 | -98.7 | -91.2 | |
| oeeee + C6H10O7 + O2 -> H2O + ooeOe-R | 3 | -100.6 | -91.2 | |
| oeeee + C6H10O7 + O2 -> H2O + ooeOe-S | 3 | -99.6 | -91.8 | |
| oeeee + C6H10O7 + 1/2O2 -> H2O + H2 + Oyeee-R | 3 | -39.0 | -40.6 | o1, o3, g |
| oeeee + C6H10O7 + 1/2O2 -> H2O + H2 + Oyeee-S | 3 | -38.8 | -40.7 | |
| oeeee + C6H10O7 + 1/2O2 -> H2O + H2 + Oeeye-R | 3 | -42.2 | -49.4 | |
| oeeee + C6H10O7 + 1/2O2 -> H2O + H2 + Oeeye-S | 3 | -41.7 | -48.8 | |
| oeeee + 2H3O+ + SO4-2 + O2 -> 3H2O + soeoe-R | 3 | -120.1 | -513.5 | o1 x 2, s |
| oeeee + 2H3O+ + SO4-2 + O2 -> 3H2O + soeoe-S | 3 | -120.6 | -514.0 | |
| oeeee + 3/2O2 -> H2 + oxoee-R | 4 | -171.7 | -158.0 | o1 x 2, o3, o2 |
| oeeee + 3/2O2 -> H2 + oxoee-S | 4 | -169.7 | -156.7 | |
| oeeee + O2 -> 2H2 + oyeye-RR | 4 | -84.3 | -81.5 | o1 x 2, o3 x 2 |
| oeeee + O2 -> 2H2 + oyeye-RS | 4 | -85.2 | -81.5 | |
| oeeee + O2 -> 2H2 + oyeye-SR | 4 | -82.5 | -80.5 | |
| oeeee + O2 -> 2H2 + oyeye-SS | 4 | -82.9 | -79.7 | |
| oeeee + O2 -> 2H2 + oeeyy | 4 | -81.9 | -81.0 | |
| oeeee + O2 -> H2 + H2O + oyoee-4D-R | 4 | -82.5 | -75.9 | o1 x 2, o3, 4D |
| oeeee + O2 -> H2 + H2O + oyoee-4D-S | 4 | -81.4 | -75.0 | |
| oeeee + O2 -> H2 + ooeye-5A-RR | 4 | -91.7 | -85.4 | o1 x 2, o3, 5A |
| oeeee + O2 -> H2 + ooeye-5A-RS | 4 | -90.8 | -84.9 | |

| | | | | |
|---|---|---|---|---|
| oeeee + O2 -> H2 + ooeye-5A-SR | 4 | -92.2 | -87.0 | |
| oeeee + O2 -> H2 + ooeye-5A-SS | 4 | -93.7 | -88.1 | |
| oeeee + O2 -> H2 + oeoye-5A-RR | 4 | -95.4 | -91.6 | |
| oeeee + O2 -> H2 + oeoye-5A-RS | 4 | -95.3 | -93.4 | |
| oeeee + O2 -> H2 + oeoye-5A-SR | 4 | -95.1 | -92.2 | |
| oeeee + O2 -> H2 + oeoye-5A-SS | 4 | -94.9 | -90.5 | |
| oeeee + O2 -> H2 + oeeyo-5A-RR | 4 | -92.0 | -88.9 | |
| oeeee + O2 -> H2 + oeeyo-5A-RS | 4 | -94.3 | -88.1 | |
| oeeee + O2 -> H2 + oeeyo-5A-SR | 4 | -91.1 | -85.6 | |
| oeeee + O2 -> H2 + oeeyo-5A-SS | 4 | -89.6 | -85.2 | |
| oeeee + C6H10O7 + O2 -> H2O + H2 + Oxeee-R | 4 | -108.4 | -103.1 | o1, o3, o2, g |
| oeeee + C6H10O7 + O2 -> H2O + H2 + Oxeee-S | 4 | -111.2 | -108.0 | |
| oeeee + C6H10O7 + O2 -> H2O + H2 + Oeexe-R | 4 | -112.0 | -109.6 | |
| oeeee + C6H10O7 + O2 -> H2O + H2 + Oeexe-S | 4 | -111.1 | -108.8 | |
| oeeee + C6H10O7 + O2 -> H2O + H2 + oXeee-R | 4 | -100.8 | -96.7 | |
| oeeee + C6H10O7 + O2 -> H2O + H2 + oXeee-S | 4 | -99.2 | -96.0 | |
| oeeee + C6H10O7 + O2 -> H2O + H2 + oeeXe-R | 4 | -108.4 | -106.9 | |
| oeeee + C6H10O7 + O2 -> H2O + H2 + oeeXe-S | 4 | -106.0 | -106.2 | |
| oeeee + 3/2O2 -> H2O + H2 + oxoee-4D-R | 5 | -155.4 | -142.9 | o1 x 2, o3, o2, 4D |
| oeeee + 3/2O2 -> H2O + H2 + oxoee-4D-S | 5 | -151.6 | -138.5 | |
| oeeee + C6H10O7 + O2 -> H2O + H2 + oOeye-5A-RR | 5 | -90.3 | -87.9 | o1 x 2, o3, g, 5A |
| oeeee + C6H10O7 + O2 -> H2O + H2 + oOeye-5A-RS | 5 | -89.5 | -87.0 | |
| oeeee + C6H10O7 + O2 -> H2O + H2 + oOeye-5A-SR | 5 | -90.9 | -88.6 | |
| oeeee + C6H10O7 + O2 -> H2O + H2 + oOeye-5A-SS | 5 | -91.1 | -88.4 | |
| oeeee + C6H10O7 + O2 -> H2O + H2 + ooeYe-5A-RR | 5 | -83.0 | -79.5 | |
| oeeee + C6H10O7 + O2 -> H2O + H2 + ooeYe-5A-RS | 5 | -84.2 | -84.3 | |
| oeeee + C6H10O7 + O2 -> H2O + H2 + ooeYe-5A-SR | 5 | -91.4 | -89.2 | |
| oeeee + C6H10O7 + O2 -> H2O + H2 + ooeYe-5A-SS | 5 | -85.9 | -84.4 | |
| oeeee + C6H10O7 + O2 -> H2O + 2H2 + Oyeye-RR | 5 | -76.1 | -78.2 | o1 x 2, o3 x 2, g |
| oeeee + C6H10O7 + O2 -> H2O + 2H2 + Oyeye-RS | 5 | -82.9 | -83.5 | |
| oeeee + C6H10O7 + O2 -> H2O + 2H2 + Oyeye-SR | 5 | -79.6 | -80.4 | |
| oeeee + C6H10O7 + O2 -> H2O + 2H2 + Oyeye-SS | 5 | -78.6 | -83.9 | |
| oeeee + C6H10O7 + O2 -> H2O + 2H2 + Oeeyy | 5 | -74.1 | -77.0 | |

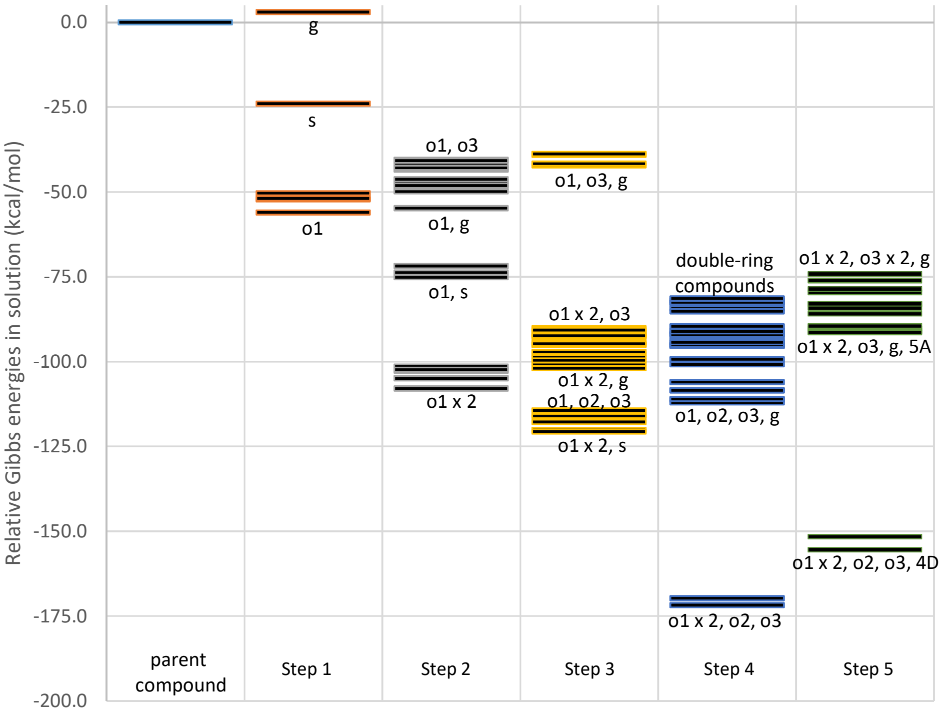

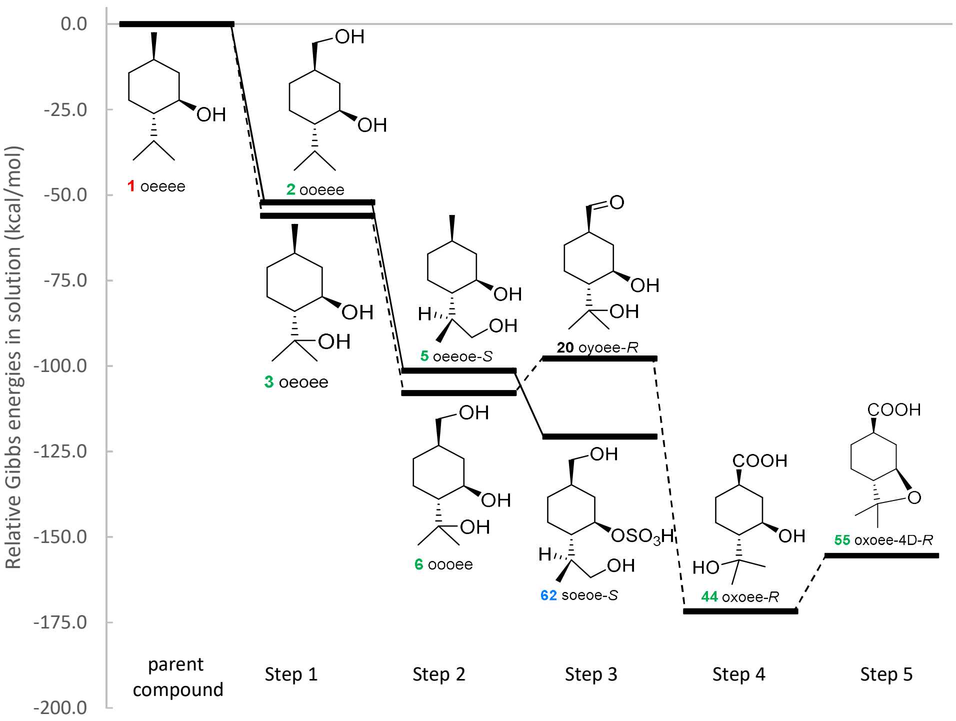

Relative Gibbs energies in solution (kcal/mol)
0.0
-25.0
-50.0
-75.0
-100.0
-125.0
-150.0
-175.0
-200.0
1 oeeee
2 ooeee
3 oeoee
5 oeeoe-S
6 oooee
20 oyoee-R
62 soeoe-S
44 oxoee-R
55 oxoee-4D-R
OH
OSO3H
COOH
HO
O
H
parent compound
Step 1
Step 2
Step 3
Step 4
Step 5

### MP2-Benchmark

| Name | Quantity | Unit | Ref |
|---|---|---|---|
| Hartree | 627.50947 | kcal mol^-1/AU | 2014 CODATA |

| Basis: 6-311++G(d,p)/Optimization in gas phase | Electronic Energy | | | | | |
|---|---|---|---|---|---|---|
| Compound | B3LYP | | MP2 | | Difference | |
| | AU | kcal/mol | AU | kcal/mol | AU | kcal/mol |
| H2 | -1.1796 | | -1.1603 | | 0.0193 | |
| H2O | -76.4585 | | -76.2749 | | 0.1836 | |
| H3O+ | -76.7312 | | -76.5501 | | 0.1811 | |
| O2 | -150.3089 | | -149.9813 | | 0.3277 | |
| O4S-2 | -699.1158 | | -697.8751 | | 1.2407 | |
| oeeee | -468.4839 | | -467.0444 | | 1.4395 | |
| oeoee | -543.7329 | | -542.1327 | | 1.6003 | |
| ooeee | -543.7214 | | -542.1178 | | 1.6037 | |
| oooee | -618.9713 | | -617.2073 | | 1.7640 | |
| ooeoe-S | -618.9605 | | -617.1930 | | 1.7675 | |
| oyoee-R | -617.7571 | | -616.0093 | | 1.7478 | |
| soeoe-S | -1242.8395 | | -1240.0348 | | 2.8047 | |
| oxoee-R | -693.0290 | | -691.1185 | | 1.9105 | |
| oxoee-4D-R | -616.5253 | | -614.7923 | | 1.7330 | |
| **Coefficient of determination** | | | | | **1.0000** | |
| **Slope** | | | | | **0.9976** | |
| **Intercept** | | | | | **0.1213** | |
| **Reaction** | **AU** | **kcal/mol** | **AU** | **kcal/mol** | **AU** | **kcal/mol** |
| oeeee + 1/2O2 -> oeoee | -0.0945 | -59.3 | -0.0976 | -61.3 | | -1.9 |
| oeeee + 1/2O2 -> ooeee | -0.0830 | -52.1 | -0.0827 | -51.9 | | 0.2 |
| oeoee + 1/2O2 -> oooee | -0.0839 | -52.6 | -0.0840 | -52.7 | | -0.1 |
| ooeee + 1/2O2 -> oooee | -0.0953 | -59.8 | -0.0989 | -62.0 | | -2.2 |
| ooeee + 1/2O2 -> ooeoe-S | -0.0846 | -53.1 | -0.0846 | -53.1 | | 0.0 |
| oooee -> oyoee-R + H2 | 0.0345 | 21.7 | 0.0377 | 23.6 | | 2.0 |
| ooeoe-S + 2H3O+ + SO4-2 -> 3H2O + soeoe-S | -0.6763 | -424.4 | -0.6912 | -433.7 | | -9.3 |
| oyoee-R + 1/2O2 -> oxoee-R | -0.1174 | -73.7 | -0.1186 | -74.4 | | -0.7 |
| oxoee-R -> oxoee-4D-R + H2O | 0.0453 | 28.4 | 0.0513 | 32.2 | | 3.8 |
| **Coefficient of determination** | | | | | | **0.9999** |
| **Slope** | | | | | | **1.0259** |
| **Intercept** | | | | | | **1.1613** |
| **Mean Absolute Error (MAE)** | | | | | | **2.25** |

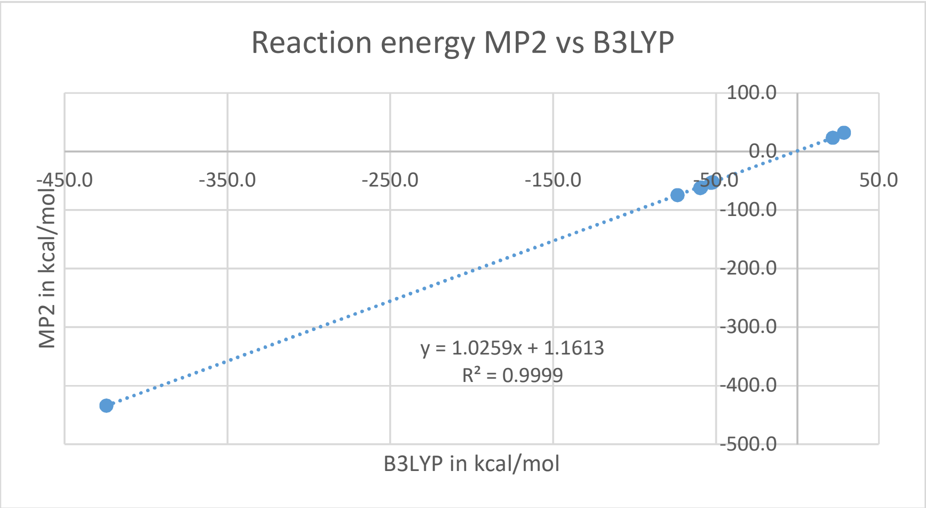
