## Supplementary material for "An *In-Silico* Investigation of Menthol Metabolism": S3 File: README.txt

DOI: 10.1371/journal.pone.0216577

6_cmpdtab.csv contains energy of all structures generated by calculations.nb in Mathematica. (raw output)
7_rxntab.csv contains energy of all stepwise reactions generated by calculations.nb in Mathematica. (raw output)
8_relstab.csv contains energy of all reactions starting from the parent compound generated by calculations.nb in Mathematica. (raw output)
calculations.xlsx is the only Microsoft Excel file in this SI that collates all key results of the paper.
 - First worksheet contains compound and reaction energy and Figure 2 of the manuscript.
 - Second worksheet contains relative stability of the 102 compounds (menthol and its metabolites) and Figures 3-4 of the manuscript.
 - Third worksheet contains MP2 calculation results of selected compounds and regression analysis for benchmarking purpose. 
calculations.pdf is a printable version of the Excel file for your convenience.  All contents are the same.
